# Consolidation-dependent behavioral integration of sequences related to mPFC neural overlap and hippocampal-cortical connectivity

**DOI:** 10.1101/2022.10.20.513126

**Authors:** Alexa Tompary, Lila Davachi

## Abstract

Systems consolidation theories propose two mechanisms that enable the behavioral integration of related memories: coordinated reactivation between hippocampus and cortex, and the emergence of cortical traces that reflect overlap across memories. However, there is limited empirical evidence that links these mechanisms to the emergence of behavioral integration over time. In two experiments, participants implicitly encoded sequences of objects with overlapping structure. Assessment of behavioral integration showed that response times during a recognition task reflected behavioral priming between objects that never occurred together in time but belonged to overlapping sequences. This priming was consolidation-dependent and only emerged for sequences learned 24 hours prior to the test. Critically, behavioral integration was related to changes in neural pattern similarity in the medial prefrontal cortex and increases in post-learning rest connectivity between the posterior hippocampus and lateral occipital cortex. These findings suggest that memories with a shared predictive structure become behaviorally and neurally integrated through a consolidation-related restructuring of the learned sequences, providing insight into the relationship between different consolidation mechanisms that support behavioral integration.

## Introduction

There are now abundant demonstrations that after periods involving consolidation, episodic memories undergo transformations that allow individuals to integrate across overlapping experiences. Examples of enhanced behavioral integration with consolidation include transitive inference (Ellenbogen et al., 2007; Lau et al., 2010; Werchan & Gómez, 2013), the extraction of statistical regularities (Wagner et al., 2004; Durrant et al., 2011, 2011; Sweegers et al., 2014; Batterink & Paller, 2017), category learning (Djonlagic et al., 2009; Graveline & Wamsley, 2017), and more (for reviews, see: Chatburn et al., 2014; Lerner & Gluck, 2019). What neural transformations support such consolidation-dependent integration across experiences?

Theories of systems-level consolidation posit that memories initially supported by the hippocampus become distributed across the cortex, through ongoing, coordinated reactivation of prior experiences (Nadel et al., 2000; Squire et al., 1984). Through this reactivation, the memory traces that are ultimately learned by cortex are thought to be highly structured and integrated representations built up from many overlapping experiences (McClelland et al., 1995). One class of theories posits that *neural* integration is accompanied by a psychological transformation such that memories supported by cortex are more gist-like and reflect the shared aspects across multiple events (Trace Transformation Theory (TTT); Sekeres et al., 2018; Winocur et al., 2010). Thus, systems-level consolidation theories point to two neural mechanisms that could support such time-dependent *behavioral* integration of overlapping memories: (1) coordinated reactivation of memory traces between the hippocampus and cortex after learning, and (2) the emergence of cortical memory traces that reflect shared content across memories for different but overlapping events. However, evidence for this is limited and, thus, it is unclear whether and how these two mechanisms may jointly support the behavioral integration of overlapping experiences. Here, we present a new behavioral protocol to probe the implicit integration of events with overlapping content, and we use this protocol to investigate how cortical similarity and post-learning hippocampal-cortical reactivation interact in supporting consolidation-dependent behavioral integration in humans.

### Integration and Cortical Similarity

According to TTT, the *behavioral* integration of related memories should be supported by the *neural* integration of cortical memory traces that reflects their overlap. Previous work has already pointed to medial prefrontal cortex (mPFC) as a key cortical region supporting integrated memories that emerge without the aid of consolidation. For instance, damage to this region impairs key memory integration behaviors like associative inference in humans (Koscik & Tranel, 2012; Spalding et al., 2018; Warren et al., 2014) and transitive inference in rodents (DeVito et al., 2010). Furthermore, activation of this region and its connectivity with the hippocampus increases when encoding episodes containing stimuli that overlap with recently learned information (Kuhl et al., 2010; Schlichting & Preston, 2016; Zeithamova et al., 2012), providing convincing evidence of its role in memory integration. Of particular relevance to the current study, patterns of activity in mPFC become more correlated for events linked through overlapping elements (Schlichting et al., 2015; Milivojevic et al., 2015) and also carry information about the common structure across many learned relationships (Morton et al., 2020; Schuck et al., 2016). With converging evidence spanning multiple species and methods, is clear that mPFC supports the acquisition and immediate expression of integrated memories.

However, there is much less evidence in humans that the neural representations in mPFC are predictive of behavioral measures of memory integration over the course of systems-level consolidation, despite such strong behavioral evidence of consolidation-dependent behavioral integration (e.g. Chatburn et al., 2014) and the increasing involvement of mPFC in memory retrieval over time (Bonnici et al., 2012; Frankland et al., 2004; Takashima et al., 2006, 2009; Takehara-Nishiuchi & McNaughton, 2008; Woodard et al., 2007). In other words, a direct link between the two is lacking. There are some clues that such a relationship should exist. First, in rodents, neural ensembles in mPFC become more selective for common features across two different associative memories and less selective to features unique to each (Morrissey et al., 2017). Second, there is evidence for neural integration of related memories in human mPFC. Previously, we used a multi-variate pattern similarity approach to demonstrate that after a week, neural patterns during the retrieval of overlapping memories grew more correlated (Tompary & Davachi, 2017). Audrain and colleagues (2020) observed a similar increase in correlation of overlapping memories over time in mPFC, although with notable differences in their protocol and consequent findings (see Discussion). However, as neither group measured the *behavioral* integration across those overlapping memories, the link between these time-dependent neural transformations and behavioral integration remains unknown. Thus, despite its theorized role in the expression of behaviorally integrated memories over time (Preston & Eichenbaum, 2013), to our knowledge, no other human research has examined how consolidation may organize memory traces in mPFC to support their behavioral integration.

### Integration and Coordinated Reactivation

Another fundamental component of systems-level consolidation theories is the coordinated reactivation of memory traces between the hippocampus and cortex after learning. One way to operationalize such post-learning coordination is by measuring how the correlation of their BOLD signals during resting-state fMRI changes after encoding new memoranda (Tambini et al., 2010; van Kesteren et al., 2010; Tompary et al., 2015; de Voogd et al., 2016; Gruber et al., 2016; Schlichting & Preston, 2016; Murty et al., 2017, 2018; Liu et al., 2018; Tambini & D’Esposito, 2020; Audrain, 2020). The vast majority of studies investigating post-learning rest connectivity find that the magnitude of experience-dependent change relates to subsequent memory accuracy, with the exception of one demonstration that post-learning connectivity relates to explicit associative inference (Schlichting & Preston, 2016). To our knowledge, this is the only finding that relates post-learning rest connectivity to a measure of behavioral integration rather than memory accuracy of learned items, but it does not examine consolidation-related changes as the integration test was administered a few minutes after learning. Furthermore, while a handful of studies reveal relationships between post-learning rest connectivity and the neural integration of overlapping memories (Audrain, 2020; Tompary et al., 2015), none to our knowledge has examined how these neural measures, jointly or separately, support behavioral integration. This leaves open the fundamental question of whether post-encoding reactivation supports the neural *transformation* of memories that renders them more integrated with other, similar memories, as predicted by TTT.

Which cortical areas, and which regions of the hippocampus, might coordinate their processing to support behavioral integration? Beginning with cortex, the earliest demonstration of post-learning connectivity implicate the involvement of category-selective cortex rather than prefrontal regions (Tambini et al., 2010). Numerous findings since then have shown that post-learning cortical connectivity with the hippocampus is governed by the category of the encoded memoranda (Collins & Dickerson, 2018; Keller & Just, 2016; Murty et al., 2017; Schlichting & Preston, 2014; Vilberg & Davachi, 2013). Because our stimuli are common objects, we focused on lateral occipital cortex, a region causally linked to the formation of object memories (Tambini & D’Esposito, 2020). Turning to the hippocampus, while the bulk of post-learning connectivity investigations treat the hippocampus as a unitary structure, there is some work that investigates differential connectivity with cortical regions along its long axis. Here, there are observations that post-learning connectivity between posterior hippocampus and category-selective cortical regions relate to later memory (Murty et al., 2017; Tambini et al., 2010), whereas connectivity between anterior hippocampus and FFA has been shown to relate to memory for faces (Liu et al., 2018) and to associative inference across stimuli including faces (Schlichting & Preston, 2016). Connectivity with anterior or posterior hippocampus also depends on whether memory is rewarded, with memory performance for low-reward items correlated with connectivity between posterior hippocampus and category-selective cortex (Murty et al., 2017). Taken together, the factors that may explain differential connectivity patterns along the long-axis of the hippocampus remain difficult to tease apart: possibilities include the form of behavior tested (memory for discrete events versus integration across them), the cortical regions selective to the content of encoded material, or the intrinsic or extrinsic motivation for consolidation. The present study did not implement a reward manipulation, so we chose to focus on posterior hippocampus, mirroring past experiments (Murty et al., 2017; Tambini et al., 2010). It is worth noting that we considered this choice to be exploratory.

### Integration of sequential regularities

Rather than employing a more traditional explicit inference task, we generated sequences ending in overlapping and distinct objects (i.e. paradigmatic relations; Luo & Zhao, 2018; McNeill, 1963). We made this decision for several reasons: (1) the learning of statistical regularities relies on the hippocampus and benefits from consolidation processes (Durrant et al., 2011, 2013; Turk-Browne et al., 2009; Schapiro et al., 2012, 2014), in line with predictions from TTT; (2) a statistical learning protocol enabled us to develop a behavioral task that relied on response times, an ideal readout of memory integration through cortical similarity and hippocampal-cortical connectivity.

In two experiments, participants viewed sequences of three objects in an incidental encoding task. Each sequence was constructed such that the first two objects always appeared consecutively (A and B), and the third object alternated between two objects (C_1_ and C_2_; Figure 1C). We reasoned that after repeated exposures to these sequences, participants would come to anticipate the presentation of two different objects (C_1_ and C_2_) upon encountering the beginning of each sequence (A and B). This expectation would give rise to a link between items C_1_ and C_2_ strictly based on their shared antecedents. Thus, although C_1_ and C_2_ never co-occurred, we predicted that the two objects would become associated through their overlapping preceding A and B objects. We tested this prediction with a novel recognition priming task, which was developed to assess the implicit behavioral integration of overlapping sequences while minimizing intentional retrieval strategies at test. An explicit memory test was also conducted for comparison with the implicit test and to link to prior work. Including both implicit and explicit memory tests also enabled us to explore changes in memory behaviors that may be underpinned by different memory processes (Henke, 2010).

**Figure 1.**
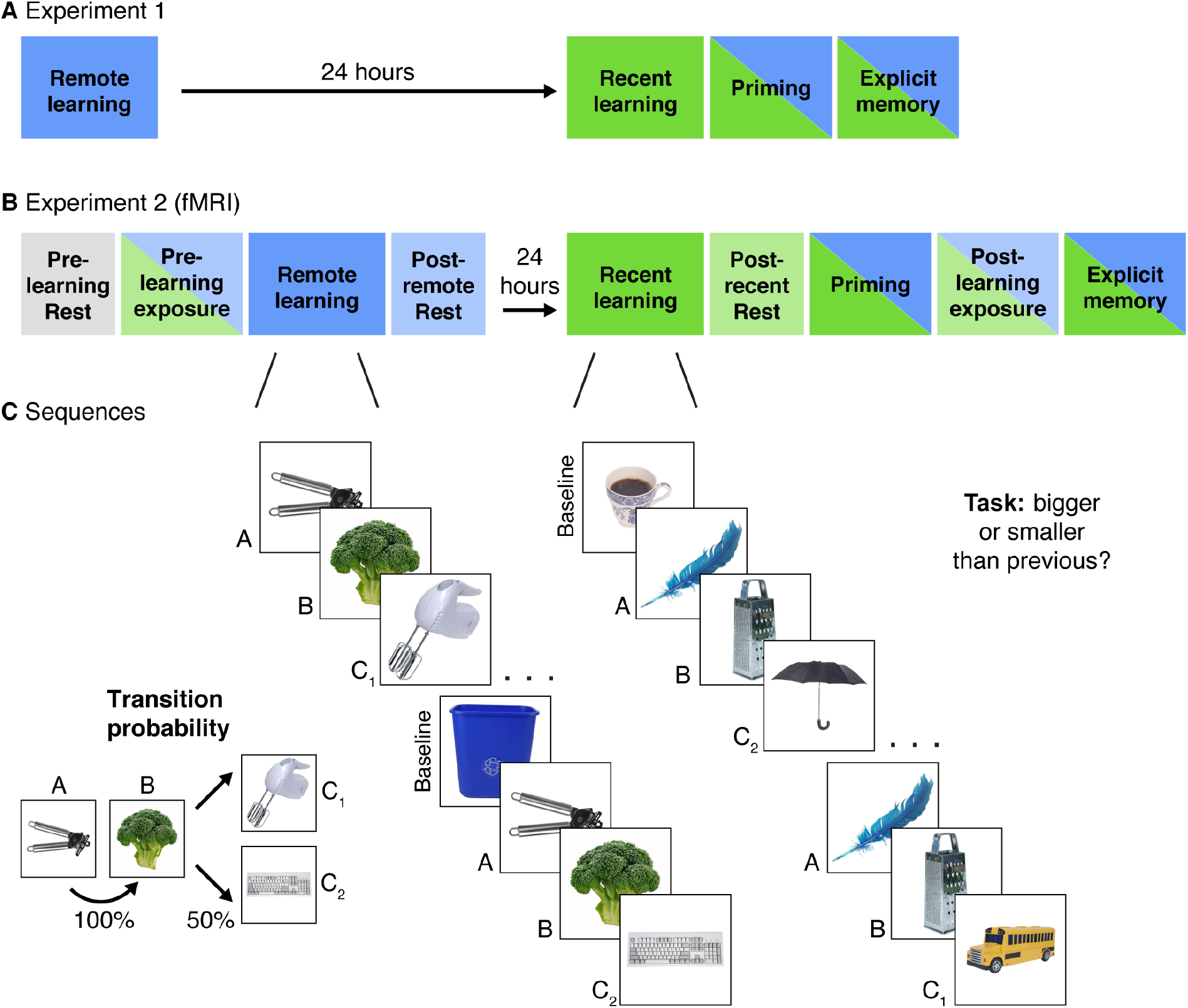
**(A)** Experiment 1 design. Participants completed two learning sessions separated by 24 hours (blue = remote, green = recent learning). **(B)** Experiment 2 design. The timing, instructions and procedures for the learning and priming tasks were identical to Experiment 1, with a modified but conceptually similar explicit memory task. Participants also completed pre- and post-learning exposure scans where all objects from both learning sessions were presented in a randomly intermixed order. Three rest scans were added: a baseline pre-learning scan at the start of Day 1, a post-remote learning scan on Day 1, and a post-recent learning scan on Day 2. **(C)** In each learning session, participants performed a cover task as they viewed objects embedded in triplets. All triplets were composed of two images that always appeared back-to-back (A and B) and two images that alternated following the AB pair (C_1_ and C_2_) with equal probability. Baseline objects were randomly inserted between triplets. Separate images sets were presented in each learning session. After the recent learning session, participants completed recognition priming and explicit memory tests over all objects intermixed from the two learning sessions (see Figure 2 for details of each test).

In Experiment 2, the same tasks were completed while participants underwent fMRI. We investigated how the observed pattern of behavioral integration related to pattern similarity in mPFC as well as changes in rest connectivity between posterior hippocampus and LOC. We predicted that neural patterns in mPFC would become more correlated for objects from overlapping sequences, but only after a period of consolidation. We further predicted that correlations between the hippocampus and LOC in rest connectivity would increase after learning, and that both measures would relate to the extent of behavioral priming participants exhibited after a delay. Finally, we examined the relationship between mPFC similarity and rest connectivity and their association with behavioral priming.

## Results

### Learning

During learning, participants were instructed to evaluate if the currently viewed object was bigger or smaller than the previous one. We used participants’ response times to assess learning of the statistical structure embedded in the learning task and to confirm that sequence learning did not differ across the recent and remote learning sessions. Because B predictably followed A and either C_1_ or C_2_ predictably followed B, we reasoned that responses to B and C objects would be facilitated after repeated exposures to their sequential structure, resulting in decreased RTs relative to A and baseline objects, which were unpredictable. As an index of learning, we computed the response times (RTs) of participant’s size judgments as a function of whether the response of each object could be predicted by the presentation of the prior one.

#### Experiment 1

We computed the median RT of all A and baseline objects (unpredictable) and the median RT of all B and C objects (predictable), for each participant’s 16 runs during each learning session. These values were entered into a mixed effects linear model with repetition (continuous: 1 – 16), learning session (discrete: recent, remote), predictability (discrete: predicted, unpredicted), and their interactions as independent variables. This model revealed an effect of repetition (F_(1, 4448)_ = 1070.17, p < .001), reflecting faster responses times as learning progressed, and an effect of predictability (F_(1, 232.4)_ = 4.48, p = .04) such that participants responded more quickly to predictable objects over unpredictable ones. These two effects were qualified by an interaction (F_(1, 4448)_ = 104.73, p < .001), reflecting a steeper drop in response times over learning for predictable objects relative to unpredictable ones (Supplemental Figure 1A). This confirms that participants learned the structure of the sequences, as by the end of learning, responses were more facilitated by predictable objects over unpredictable ones.

The same model also revealed differences between the recent and remote learning sessions, with an effect of learning session (F_(1, 119.9)_ = 17.63, p < .001) and an interaction between repetition and learning session (F_(1, 4448)_ = 4.57, p = .03). These effects were driven by overall slower RTs and a steeper change in RTs across learning in the remote session relative to the recent one, suggesting that responses in the recent session were in part driven by acclimation to the task. Critically, there was no interaction between learning session and predictability or between learning session, predictability, and repetition (both F’s < 2.10, both p’s > .14) suggesting that participants became sensitive to the sequence structure and were faster to respond to predictable objects over unpredictable ones during both sessions.

Average accuracy on the size judgment task was high (92.6%; SD = 12.5%). Changes in accuracy over learning were analyzed with the same mixed effects linear model with repetition (continuous: 1 – 16), learning session (discrete: recent, remote), predictability (discrete: predicted, unpredicted), and their interactions as independent variables (Supplemental Figure 1B). On average, accuracy did not reliably differ by learning session, repetition, or predictability (all F’s < 2.57, all p’s > .11). However, this model revealed several interactions: between repetition and predictability (F_(1, 4448)_ = 7.82, p = .005), with higher accuracy for predictable objects over unpredictable objects towards the end of learning relative to the beginning; between learning session and predictability (F_(1, 4448)_ = 4.07, p = 0.04), with a more pronounced difference in accuracy by predictability in the recent session over the remote; and between repetition and learning session (F_(1, 4448)_ = 9.72, p = .002), with a more pronounced increase in accuracy over learning in the remote session relative to the recent one. Taken together, this suggests that although size comparisons were overall very accurate, responses were influenced by the predictability of the objects and the order of the learning sessions.

#### Experiment 2

Participants exhibited the same changes in response times as in Experiment 1 (Supplemental Figure 1A). Participants became faster across repetitions (F_(1, 1459)_ = 354.21, p < 0.001) and for predicted relative to unpredictable objects (F_(1, 43.06)_ = 19.00, p < .001). Critically, the interaction between predictability and repetition was replicated (F_(1, 1459)_ = 34.66, p < .001) with more facilitated responses across repetitions for predictable relative to unpredictable objects. The same effects of learning session were present as well: faster response times during recent relative to remote sessions (F_(1, 40.69)_ = 22.66, p < .001), and a larger facilitation of RTs across repetitions in the remote session over the recent session (F_(1, 1459)_ = 32.16, p < .001). As in Experiment 1, there were no other interactions with learning session (both F’s < 2.65, both p’s > .10), suggesting that learning was equivalent across the two sessions.

Average accuracy on the size judgment task was consistently high (95.3%; SD = 7.4%) and changes in accuracy over learning were largely similar to those observed in Experiment 1 (Supplemental Figure 1B). Specifically, accuracy was modulated by an interaction between repetition and predictability (F_(1, 1459)_ = 8.61, p = .003), reflecting higher accuracy for predictable over unpredictable objects towards the end of learning, and an interaction between repetition and learning session (F_(1, 1459)_ = 5.66, p = .02), with a more pronounced increase in accuracy over learning in the remote session relative to the recent one. There were no reliable effects of learning session or predictability and no other interactions (all F’s < .99, all p’s > 32), with the exception of an effect of repetition (F_(1, 1459)_ = 713, p = .008) not observed in Experiment 1.

### Recognition

As a reminder, although C_1_ and C_2_ were never experienced together in time, we predicted that the two objects would become associated through their overlapping preceding A and B objects, and that this across-sequence association would become strengthened over time. To test this prediction, we developed a novel recognition priming task in which participants viewed all objects from both learning sessions intermixed with novel foils and endorsed each as ‘old’ or ‘new’. Unbeknownst to the participants, the order of the objects was manipulated to use response priming as a behavioral index of participant’s implicit integration of C objects from overlapping sequences (Figure 2A, top). We developed mixed-effects models to examine differences in RTs between C_2_ objects, which were preceded by and thus primed by C_1_, and RTs for C_1_ objects, which were preceded by baseline objects and thus served as a control comparison (see Methods). To be included in this priming analysis, each pair of C objects and the preceding control object must have been correctly endorsed as ‘old’. Note that C_1_ and C_2_ are not differentiable to participants, as during learning, they both follow B an equal number of times and are presented within their sequence in a randomized and intermixed order. We assigned them with separate labels for the sole purpose of clarifying the conditions of the priming manipulation.

**Figure 2.**
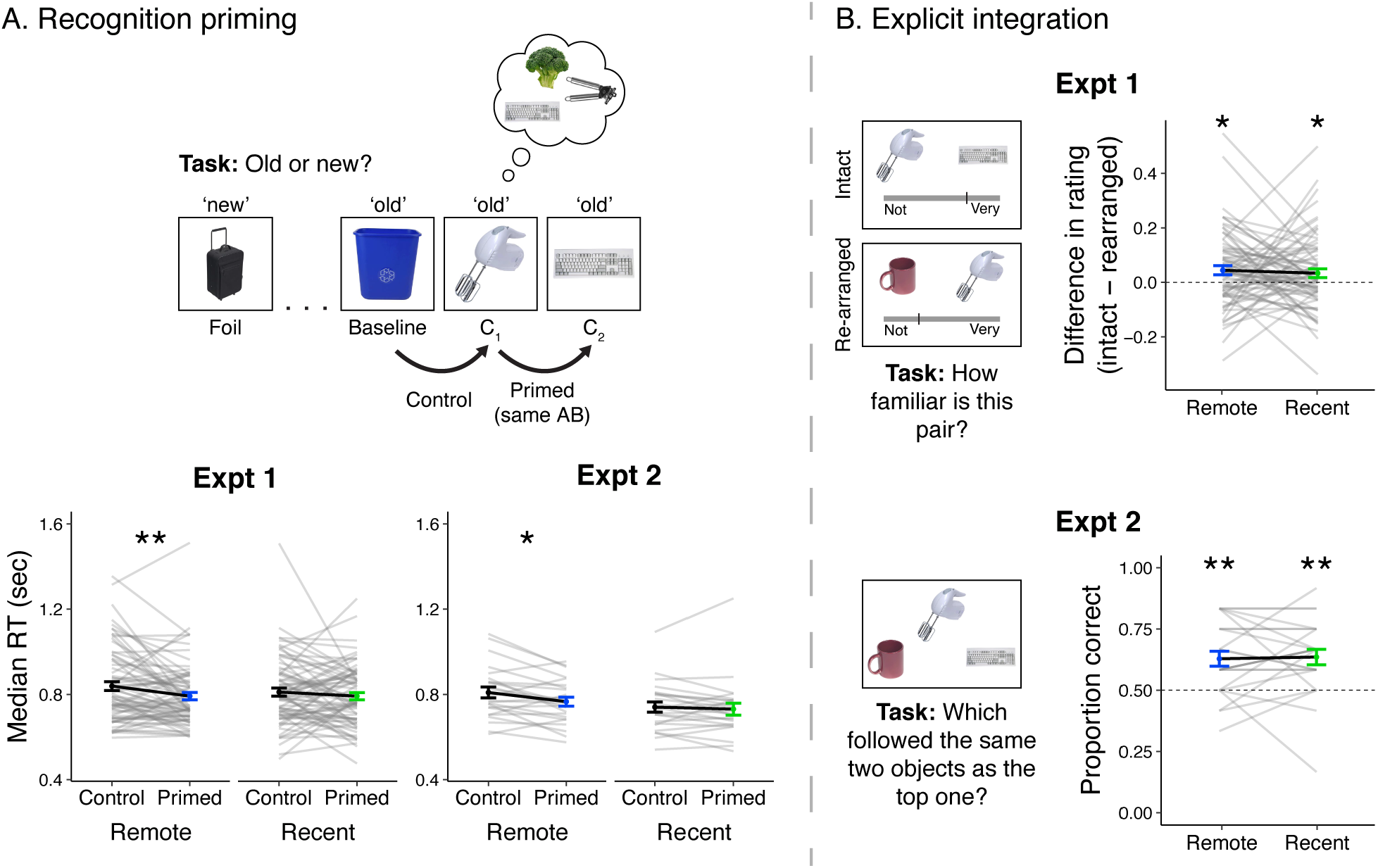
**(A, top)** Recognition priming design. Studied and new objects were presented pseudo-randomly such that each C2 object followed the C1 object from the same sequence, and the C1 object followed a baseline object from the same learning session. The behavioral integration of C1 and C2 was operationalized as the decrement in response time to C2 (preceded by C1) relative to C1 (preceded by a baseline object). **(A, bottom)** Recognition priming results. Statistics reflect results from trial-level mixed-effects models, but participants’ responses are aggregated to facilitate visualization of the effects. Black lines represent group averages. Gray lines represent median response times for each participant. Error bars indicate standard error of the mean (SEM) across participants. ** indicates p < .01; * indicates p < .05. **(B, left)** Explicit integration design. In Experiment 1, participants viewed intact and re-arranged pairs of C objects and rated their familiarity on a continuous scale from ‘Not familiar at all’ to ‘Very familiar’. All foils for re-arranged pairs were C objects from a different sequence learned in the same session. The average difference in rating for intact versus re-arranged pairs served as a measure of explicit integration across overlapping sequences. In Experiment 2, participants performed a 2AFC task with a C object as the cue and were asked to choose which object followed the same pair of two objects as the cue. Foil images were C objects from a different sequence learned in the same session. **(B, right)** Explicit integration results. Black lines represent group averages. Gray lines represent participants. Error bars indicate standard error of the mean (SEM) across participants. Statistics reflect one-sample t-tests against chance performance. ** indicates p < .01; * indicates p < .05.

#### Experiment 1

Recognition accuracy across all objects approached perfect performance, as measured by A, a non-parametric measure of sensitivity that integrates hits and false alarms and can accommodate perfect performance (remote: mean A = .967, SD = .043; recent: mean A = .967, SD = .045). A Wilcox signed-ranks test indicated no detectable difference across learning sessions (V = 683.5, p = .48).

Because each participant had different numbers of trials included, mixed effects linear models were computed with RT (continuous) as the dependent variable and order (discrete: primed, control) as a predictor, separately for recently and remotely learned objects (Figure 2A, bottom left). This decision reflected our a-priori prediction that remotely learned sequences would be more strongly integrated over recently learned ones. Indeed, we found faster response times for primed objects relative to unprimed objects for the remotely learned objects (F_(1, 212.34)_ = 7.49, p = .007) but not the recently learned objects (F_(1, 69.12)_ = 0.37, p = .54). This finding suggests that objects that are linked by overlapping preceding sequences become associated over time. To assess if these effects were different across the two learning sessions, a mixed effects model was computed with order (discrete: primed, control), learning session (discrete: recent, remote) and their interaction as predictors. This revealed an effect of order (F_(1, 70.07)_ = 5.40, p = 0.02), no reliable effect of session (F_(1, 60.45)_ = 3.40, p = .07), and no reliable interaction (F_(1, 1295)_ = 2.54, p = .11).

#### Experiment 2

Similar to Experiment 1, recognition accuracy across all objects was consistently high (remote: mean A = .977, SD = .018; recent: mean A = .983, SD = .013) with a small but reliable decrease in accuracy for remotely learned objects relative to recently learned ones (V = 12.0, p = .04).

Priming analyses for participants in Experiment 2 primarily replicated findings from Experiment 1 (Figure 2A, bottom right). As in Experiment 1, we found faster RTs for primed objects relative to control objects, for objects learned in the remote session (F_(1, 89.79)_ = 5.28, p = .02) but not the recent session (F_(1, 229.72)_ = 0.56, p = .47). However, a trial-level mixed effects model including trials from both sessions revealed an effect of order (F_(1, 71.71)_ = 4.15, p = .04), an effect of learning session (F_(1, 24.12)_ = 5.68, p = .03), and no reliable interaction (F_(1, 468.80)_ = 1.34, p = .25). Taken together, response times from both cohorts reflect a behavioral association of objects from overlapping sequences, despite never having been experienced at the same moments in time.

### Explicit integration

Our primary predictions involved the recognition priming task, as an implicit measure of association would minimize any contributions of active strategies or control processes that may contribute to integration at retrieval. However, we also included explicit tests of memory integration for comparison to other studies that report such tests (Figure 2B, left). In Experiment 1, we explored a novel task that provided us with a continuous measure of integration. In Experiment 2, we reverted to a 2-alternative forced choice (2AFC) format more commonly used to investigate neural measures underlying integration (Preston et al., 2004; Zeithamova et al., 2012).

#### Experiment 1

In this task, participants viewed pairs of C objects that either followed the same A and B objects (Intact) or were paired with different A and B objects (Rearranged) during learning. They reported the strength of their associative recognition of these pairs on a sliding scale. Behavioral integration was operationalized as the average difference in familiarity between Intact and Re-arranged pairs. A value of 0 indicates no discrimination and 1 indicates maximal discrimination between the two conditions. Accuracy was reliably above chance (remote: mean = 0.044, SD = 0.142, t_(70)_ = 2.63, p = .01; recent: mean = 0.034, SD = 0.139, t_(70)_ = 2.04, p = .046) with no reliable difference across learning sessions (t_(70)_ = 0.46, p = .65; Figure 2B, top right).

#### Experiment 2

Here, we tested explicit integration with a 2AFC task with a C object as the cue, the C object from the overlapping sequence as the target, and a C object from a different sequence as the foil. We computed participants’ average accuracy over all trials, with performance at .5 indicating chance-level performance. Average accuracy was above chance for objects learned remotely (mean = 0.628, SD = 0.149, t_(23)_ = 4.21, p < .001) and recently (mean = 0.635, SD = 0.155, t_(23)_ = 4.28, p < .001) with no difference across the two learning sessions (t_(23)_ = -0.22, p = .83; Figure 2B, bottom right). These results serve as a conceptual replication of Experiment 1, as both cohorts exhibited weak integration between C objects that did not change over time. Interestingly, these results differ from the recognition priming findings, which revealed behavioral integration only after a delay. Potential accounts for this discrepancy are considered in the Discussion.

### Experiment 2: pattern similarity

Participants in Experiment 2 completed a modified set of procedures from Experiment 1 while undergoing fMRI (Figure 1B, see Methods). Specifically, their procedure included a pre- and post-learning exposure phase in which they viewed all object images intermixed while performing a cover task, and three scans capturing periods of rest occurring pre-learning, post-recent learning, and post-remote learning. In this section, we focus on the pre- and post-learning exposure scans (Figure 3A). Presenting images before and after learning enables us to extract ‘snapshots’ of the pattern of activity evoked by each image before and after participants learned their temporal associations, and the pseudo-random trial order allowed us to statistically separate responses to individual images (Schapiro et al., 2012). Specifically, we were interested in quantifying changes in the similarity in patterns of activity evoked by C objects as a function of whether they were learned in overlapping or distinct sequences (e.g. followed by the same or different AB images) and when they were learned (either in the recent or remote learning sessions). We also examined their relationship with the recognition priming effects observed in those participants.

**Figure 3.**
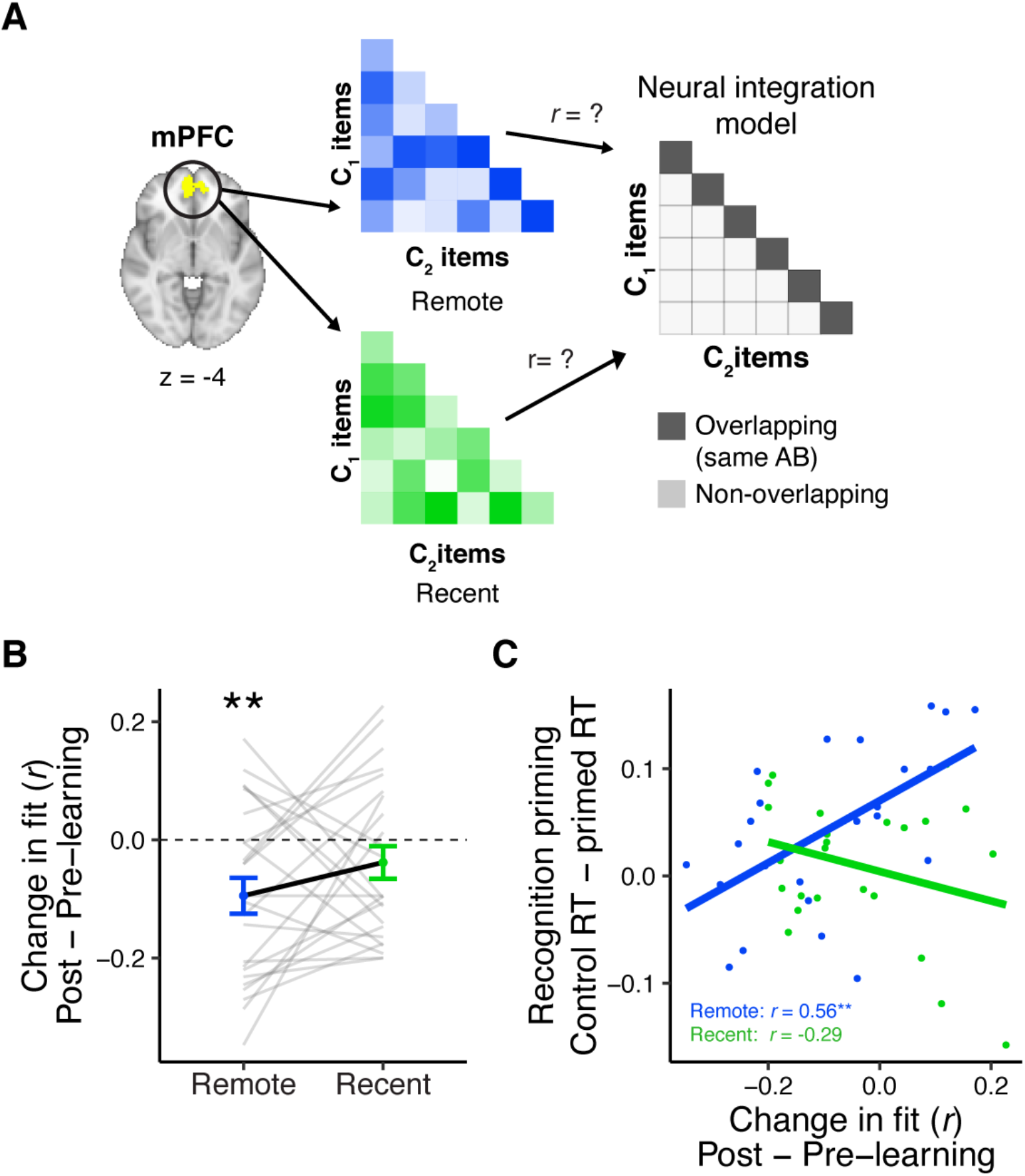
**(A)** Analysis approach for exposure scans. Vectors of multi-variate activation were extracted from an mPFC ROI for each C object and correlated with vectors for all C objects from overlapping and distinct sequences viewed in the same learning session (recent and remote). These correlation matrices were correlated with an ‘integration model’ reflecting greater similarity for C objects belonging to an overlapping sequence (i.e. objects that followed the same AB sequence). This analysis was repeated for both exposure phases and subtracted (post-minus pre-learning exposure) to create change scores. **(B)** Average change in fit to the integration model (post-learning minus pre-learning) in mPFC, separately for recently and remotely learned sequences. Values < 0 indicate a worse fit to the model after learning, meaning differentiation of C objects from overlapping sequences relative to C objects from different sequences. Gray lines indicate participants. Black line indicates group average. Error bars reflect SEM. ** indicates p < .01 relative to 0. **(C)** Correlation between the average change in the fit to the integration model in mPFC and average recognition priming across participants, separately for recent and remote learning. Dots indicate participants. Lines indicate line of best fit.

#### Learning-related and time-dependent changes in similarity

We quantified changes in similarity between all C objects by extracting patterns of activation for each C object and compiling the correlation between all patterns into a matrix, separately for sequences learned recently or remotely and separately for the pre- and post-exposure phases. We correlated these four matrices with a model matrix postulating greater similarity between C objects that followed the same AB sequence relative to those that followed different AB objects (‘neural integration model’; Figure 3B). We then subtracted the pre-learning fit from the post-learning fit for recently and remotely learned objects, resulting in a change score where values greater than 0 indicate that C objects following the same AB sequence became more similar to each other relative to C objects from other sequences. Change values less than 0 indicate that C objects became more differentiated. As a reminder, the pre-learning exposure phase took place immediately before remote learning on Day 1, and the post-learning exposure phase took place after recent learning on Day 2. This design allows us to dissociate changes in similarity that emerge from sequence learning alone (recent learning), versus from learning followed by a period of consolidation (remote learning).

We applied this analysis to multi-voxel patterns in a functionally defined mPFC region from the learning scans (see Methods). This region and others exhibited greater univariate activation for unpredictable objects (i.e. A and baseline objects) relative to predictable objects (e.g. B and C objects), when considering all learning scans in both sessions (Supplemental Figure 2A). Surprisingly, in mPFC, the change in fit between the correlation matrix reflecting neural similarity and the model was significantly negative for the remotely learned objects (t_(23)_ = -3.06, p = .006; Figure 3B). A negative change score reflects a *decrease* in fit between the multi-voxel patterns and the integration model in the post-learning relative to pre-learning snapshots, which means that over a combined period of learning and consolidation, voxels in mPFC *differentiated* objects that appeared in overlapping sequences, contrary to our prediction. This decrease remained when accounting for variation in BOLD signal (see Supplementary Information). There was no reliable change in fit to the neural integration model for recently learned objects (t_(23)_ = - 1.37, p = .18), however there was no reliable difference in the extent of the change in fit between recent and remote learning (t_(23)_ = -1.42, p = .17).

#### Relationship between mPFC similarity and priming

Next, we asked whether changes in mPFC pattern similarity tracked the recognition priming observed for objects whose sequences were learned 24 hours prior. To do this, we quantified the priming effect for each participant as the median of the difference in RTs between responses to primed and unprimed C objects, where a greater difference score indicates a stronger priming effect. We then correlated this score with participants’ change in fit to the neural integration model from the pre- to post-learning exposure phases. We computed this correlation separately for the objects learned recently and remotely (Figure 3C). We found that participants with *increased similarity* amongst C pairs from overlapping sequences exhibited a stronger priming effect, but only for pairs learned remotely (*r*_(22)_ = .57, p = 0.004). This relationship was not detectable for pairs learned recently (*r*_(22)_ = -.30, p = 0.15) and the difference in the two correlations was significant (z = 3.08, p < .001). This suggests that across participants, the extent of neural integration in mPFC reflects the extent of behavioral integration across overlapping sequences, but this association only emerges after a period involving consolidation.

#### Changes in similarity in other regions

Prior work has also observed neural integration in category-selective cortical regions immediately after learning (Richter et al., 2016; Tompary & Davachi, 2017; Wing et al., 2020). Using the same neural integration model as with mPFC, we analyzed patterns of activity in LOC, a region sensitive to object stimuli. We found that C objects belonging to overlapping sequences grew more similar to each other after recent learning (t_(23)_ = 3.10, p = .005) but this effect was not evident for the remotely learned items (t_(23)_ = 0.38, p = .71), with no reliable difference across the two (t_(23)_ = 1.98, p = .06). Unlike in mPFC, the extent of this change did not relate to behavioral priming across participants (both *r*’s < .3, both p’s > .15). No hippocampal ROIs exhibited changes in their fit to the neural integration model (all t’s < 0.94, all p’s > .35).

Taken together, we find that although on average, objects who appeared separately in time, but shared overlapping antecedents, are integrated in LOC immediately after learning but are differentiated in mPFC after 24 hours. However, despite this differentiation at the group-level, participants with stronger neural integration in mPFC exhibited facilitated behavioral integration as reflected by stronger recognition priming.

### AB integration and similarity

While the primary focus of this experiment was investigating behavioral and neural integration *across* sequences with overlapping regularities, we also included tests of memory for AB pairs, to confirm that participants were able to learn those components of each sequence. We observed consistently high explicit memory for the pairs across both experiments, and implicit integration after a delay in Experiment 1 only (Supplemental Figure 5). We also designed Experiment 2 to replicate findings of integration *within* sequences by assessing the similarity of A and B objects from overlapping versus different sequences (Supplemental Figure 6). We found that AB pairs were more neurally integrated in the anterior aspect of the hippocampus as well as in lateral occipital cortex, but only for pairs learned recently and not remotely. This is a conceptual replication of prior work showing neural integration of temporally co-occurring shapes in the hippocampus immediately after learning (Schapiro et al., 2012).

### Experiment 2: resting state connectivity

Having established a behavioral measure representing time-dependent integration across events and linking that behavior to a neural measure of integration, we next were interested in understanding whether these emerging signals were supported by active consolidation processes. As mentioned in the introduction, here we focus on changes in resting state connectivity between the hippocampus and category-selective cortex. Rest connectivity is operationalized here as the Pearson correlation between the time courses of two ROIs (Figure 4A).

**Figure 4.**
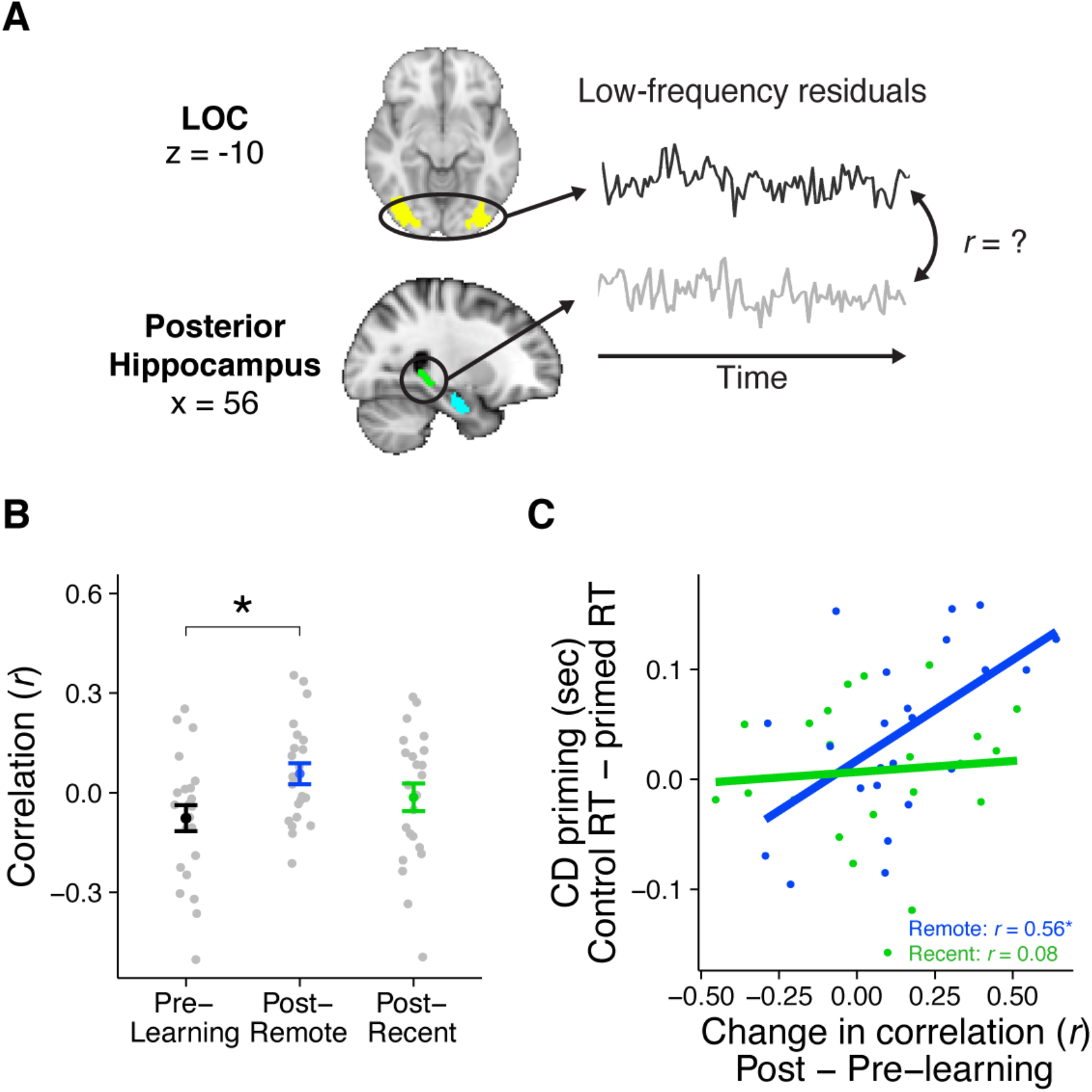
**(A)** Analytic approach for rest scans. Rest scans were preprocessed, stripped of nuisance signals, and band-pass filtered. The mean residual signal was extracted from the posterior hippocampus and LOC for each volume of each scan. Rest connectivity was measured by correlating their mean time courses. **(B)** Average rest connectivity between LOC and posterior hippocampus. Gray dots indicate participants. Black, blue, and green dots indicate group averages. Error bars reflect SEM. * indicates p < .05. **(C)** Correlation between the change in LOC - posterior hippocampal rest connectivity and average recognition priming across participants, separately for recent and remote learning. Dots indicate participants. Lines indicate best fit.

#### Learning-related changes in rest connectivity

To first examine broad changes resulting from sequence learning, we collapsed across both recent and remote learning and found a reliable increase in rest connectivity between posterior hippocampus and LOC (t_(22)_ = 2.26, p = .03). This increase was driven primarily by an increase in connectivity after remote learning (occurring on Day 1) (t_(22)_ = 2.70, p = .01) rather than after recent learning (occurring on Day 2) (t_(22)_ = 1.11, p = .28), although the extent of these increases were not reliably different from each other (t_(22)_ = 1.23, p = .23). Notably, Day 1 of the experiment (comprising pre-learning rest, remote learning, and post-remote learning rest) is most analogous to prior designs investigating learning-related changes in connectivity, so the observation that hippocampal-LOC connectivity increases reliably after remote learning is a conceptual replication of those findings (Murty et al., 2017; Tambini et al., 2010). This finding was selective to aspects of the posterior hippocampus, as there was no analogous change in rest connectivity between anterior hippocampus and LOC (t_(22)_ = 0.23, p = .82).

Since mPFC exhibited changes in neural patterns that reflect learning and consolidation of the sequences, we also asked whether rest connectivity with this region changed after learning. Surprisingly, there was an increase in rest connectivity between LOC and mPFC (t_(22)_ = 2.25, p = .03). This increase was driven primarily by an increase after recent learning, on Day 2, (t_(22)_ = 2.31, p = .03) but not remote learning, which took place on Day 1 (t_(22)_ = 1.18, p = .25), with no reliable difference in their extent (t_(22)_ = -1.00, p = .33; Supplemental Figure 4A). There was no reliable change in rest connectivity between mPFC and either anterior or posterior hippocampus (both t’s < 1.23, both p’s > .23), suggesting a selective role of sensory cortex in post-learning processing of the sequences consistent with prior work (Murty et al., 2017; Tambini et al., 2010).

#### Relationship between rest connectivity and priming

As past work has observed that learning-related changes in rest connectivity are related to later memory for the preceding learned items (Murty et al., 2017; Tambini et al., 2010; Tompary et al., 2015), we next investigated whether changes in rest connectivity between posterior hippocampus and LOC related to our measures of integration as evidenced by recognition priming (Figure 4C). To do this, we conducted two across-participant correlations between the change in rest connectivity (post-learning minus pre-learning) and median recognition priming of sequences: one from the remote learning session, and another from the recent learning session. We found that participants with a greater change in posterior hippocampal-LOC connectivity exhibited larger priming effects only for sequences from the remote learning session (r_(21)_ = .56, p = .005) and not the recent learning session (r_(21)_ = .09, p = .69). This relationship between connectivity and behavior was not significantly different across the two sessions (z = 1.68, p = .09).

We conducted several control analyses to investigate the specificity of the relationship between changes in rest connectivity and recognition priming from the remote session. First, rest connectivity from the recent session did not relate to remote priming, and rest connectivity from the remote session did not relate to recent priming (both *r*’s < .10, both p’s > .65), suggesting that learning-related changes in rest were related solely to the memoranda learned in the intervening session. Second, LOC rest connectivity with neither mPFC (Supplemental Figure 4B) nor anterior hippocampus related to priming from either session (all *r*’s < .23, all p’s > .28). Third, rest connectivity between LOC and posterior hippocampus did not relate to any explicit memory measures for C_1_ and C_2_ pairs at either time point (both *r*’s < -.32, both p’s > .13). Together, these observations demonstrate that the long-term implicit integration across overlapping sequences is selectively related to immediate post-learning rest connectivity between the posterior hippocampus and LOC.

### Experiment 2: Relationship between neural integration and rest connectivity

So far, we have reported that priming of overlapping sequences is related both to changes in their neural similarity in mPFC and increases in rest connectivity between posterior hippocampus and LOC. In the next section, we aimed to examine the relationship between these two neural measures and test whether they contributed unique variance in the extent of recognition priming observed across participants.

#### Relationship between similarity and rest

The relationship between mPFC similarity and rest connectivity was computed by correlating the two measures across participants separately for the recent and remote learning sessions (Figure 5A). We found that participants with a greater increase in mPFC similarity of overlapping sequences also exhibited a greater increase in rest connectivity immediately post-learning, but only for objects from the remote session (r_(21)_ = .42, p = .048) and not the recent session (r_(21)_ = - .18, p = .40). Furthermore, this relationship was reliably stronger for remote session over the recent session (z = 2.04, p = .04).

**Figure 5.**
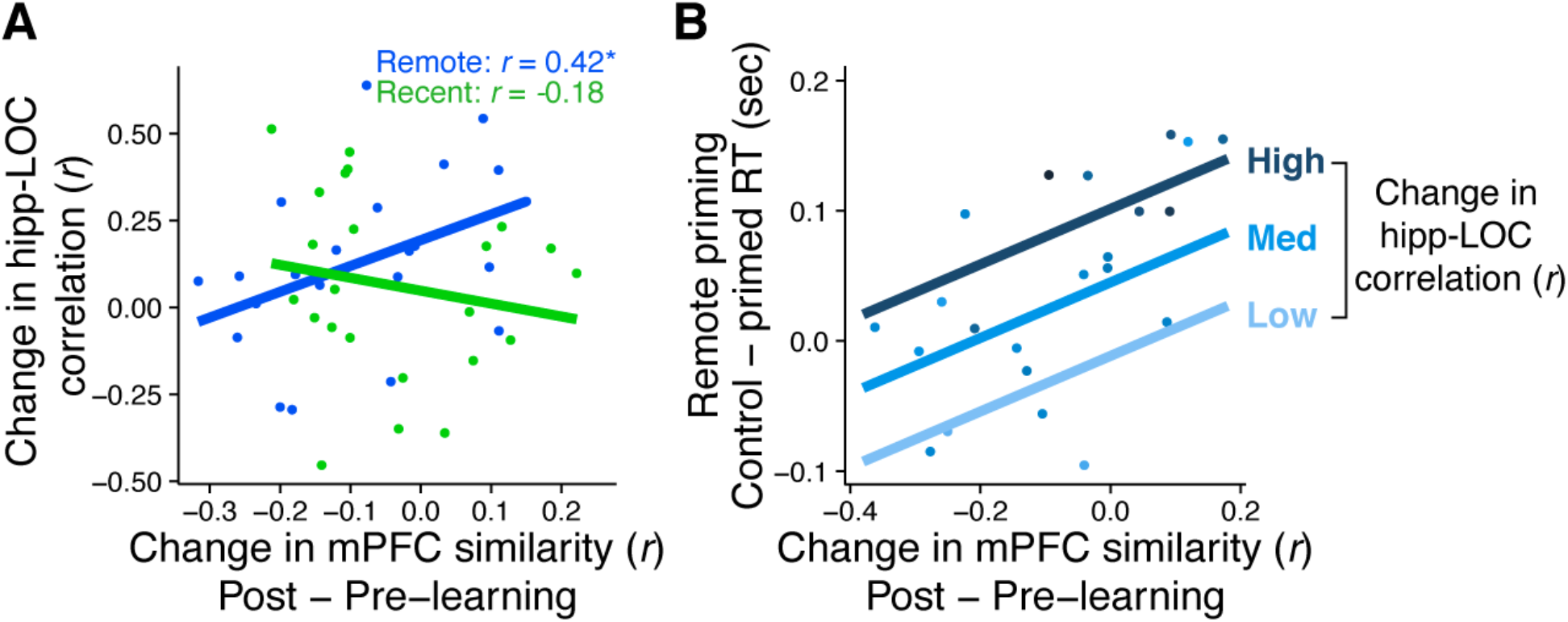
**(A)** Correlation between the change in mPFC similarity and change in rest connectivity between posterior hippocampus and LOC across participants. Dots represent participants and lines represent model fit. * indicates p < .05. **(B)** Visualization of multiple linear regression predicting recognition priming for the remotely learned sequences. Each dot is a participant. Color indicates the magnitude of the change in rest connectivity between posterior hippocampus and LOC. Lines represent fits of the model at different magnitudes of rest connectivity.

#### Neural measures related to priming

Since the neural measures of mPFC similarity and LOC-hippocampal rest connectivity were correlated for the remotely learned sequences, we next assessed whether the two measures explained unique or shared variance in their relationship with recognition priming. Focusing on data from the remote learning session, we computed a multiple regression with recognition priming (i.e., the median differences in primed vs unprimed response times) as the dependent variable, and two predictors: the change in rest connectivity and the average change in mPFC similarity. We then compared this full model against two models, each missing either mPFC similarity or rest connectivity as a predictor. These comparisons revealed unique contributions of both change in mPFC similarity (F_(1, 21)_ = 5.63, p = .03) and change in posterior hippocampal-LOC rest connectivity (F_(1, 21)_ = 4.55, p = .045) in explaining recognition priming across participants. Figure 5B visualizes the results of this model, showing participants’ relationship between their recognition priming of the remotely learned objects and their change in mPFC similarity. This relationship is overlaid with lines of best fit that represent the relationship between mPFC similarity and priming given three magnitudes of hippocampal-LOC connectivity. This suggests that both measures are *positively and uniquely* associated with participants’ time-dependent integration across overlapping sequences.

## Discussion

Across two experiments, we investigated how cortical representations and post-learning reactivation influenced the behavioral integration of memories with overlapping predictive structure. We manipulated overlap in sequences of triplets such that objects that never occurred together in time shared antecedents (i.e., both were predicted by the appearance of the same pair of objects). First, we found that participants’ response times reflected increased association of objects with shared temporal structure, but only 24 hours after learning the sequences. This delay-dependent behavioral measure of integration was replicated in a cohort that underwent fMRI. In this cohort, we found that cortical neural similarity was shaped by sequence learning in a delay-dependent manner: patterns of activity in LOC reflected the immediate association of objects from overlapping sequences, while mPFC differentiated objects from overlapping sequences learned 24 hours prior. At the same time, learning of the sequences increased post-learning connectivity between the posterior hippocampus and LOC. Critically, after a 24-hour delay, changes in mPFC similarity and changes in hippocampal-LOC rest connectivity were correlated across participants, and both measures explained unique variance in the extent of recognition priming across participants. We interpret these findings as evidence that both coordinated reactivation and cortical learning are markers of consolidation that contribute to the behavioral integration of overlapping memories over time. Below, we discuss each of these findings through the lens of systems-consolidation theories and in the context of prior empirical work.

### Integration of overlapping regularities

As a reminder, we observed implicit integration across overlapping sequences in a recognition priming protocol, only after a 24-hour period, suggesting that systems-level consolidation processes can enhance behavioral integration across memories with overlapping content. This finding adds to the literature showing consolidation-dependent memory integration in different tests of integration: in transitive inference (Ellenbogen et al., 2007; Lau et al., 2010; Werchan & Gómez, 2013), the extraction of statistical regularities (Wagner et al., 2004; Durrant et al., 2011, 2011; Sweegers et al., 2014; Batterink & Paller, 2017) and category learning (Djonlagic et al., 2009; Graveline & Wamsley, 2017). However, we observed weak but reliable explicit integration for the same items immediately after learning, which remained unchanged over the same 24-hour period, consistent with many observations showing immediate explicit integration across overlapping memories (Acuna, Eliassen, et al., 2002; Acuna, Sanes, et al., 2002; Ellenbogen et al., 2007; Greene et al., 2006; Heckers et al., 2004; Lau et al., 2010; Preston et al., 2004; Werchan & Gómez, 2013; Zeithamova et al., 2012). Even though both results are consistent with prior work, what could explain the discrepancy in the timing of successful implicit and explicit integration—both of which replicated in a second cohort of participants? A handful of behavioral studies find that participants perform better at explicit integration tests after sleep, but these studies differ in many factors that could explain the discrepancy with our findings: (1) instructions to explicitly encode premise pairs (Ellenbogen et al., 2007), (2) foreknowledge of the underlying structure of the stimulus associations prior to learning (Sweegers & Talamini, 2014), and (3) the use of non-temporal versus temporal structure of the regularities (Lerner & Gluck, 2019).

The paradigms that specifically investigate the integration of sequences merit special attention. There is some evidence of transitive inference across overlapping sequences (e.g. inferring AC from viewing AB and BC in a continuous stream of colored dots) (Luo and Zhao, 2018). Explicit recognition of these associations was on par with performance in our explicit integration task. However, there was no delayed test or implicit integration measure, precluding any interpretations about the role of consolidation in this form of integration. The sequences in our protocol also resemble paradigmatic relations (McNeill, 1963), which are second-order associations that do not co-occur, but instead are substitutable with each other because of their shared context (e.g. wearing ‘flip-flops’ and wearing ‘boots’). Interestingly, Yim and colleagues (2019) observe implicit learning of paradigmatic relations only when participants are instructed to attend to the stimuli (i.e. actively categorizing images rather than listening to an auditory stream while coloring) and have robustly learned the relevant premise pairs. This fits with our findings in several ways: first, our encoding task required constant comparisons between the present and previous objects, which may have heightened awareness of the temporal regularities, like in their active encoding protocol. Second, Yim and colleagues find strong evidence of explicit memory for first-order associations across all experiments, which mirrors our observations of strong explicit knowledge of AB pairs. However, one notable difference is that Yim and colleagues did not administer an implicit test for paradigmatic associations after a delay, which is the only condition where we observe integration. Why integration emerges immediately after learning in their experiment, but only after a period of consolidation in our experiment, remains an open question.

Finally, the discrepancy between our implicit and explicit tasks could be explained more broadly by differences in memory strength for the within-event and across-event associations. Results from the explicit memory test indicate much better integration of objects in the same sequence (A and B) relative to integration across sequences (C_1_ and C_2_). Importantly, A and B were viewed twice as many times as each C, as they preceded each appearance of C_1_ and C_2_. It may be that these overlearned associations are more accessible and explicit integration tests are sensitive to this information. At the same time, consolidation may promote representational changes that (1) benefit associations that are not already strengthened through sufficient learning (Schapiro et al., 2018), or (2) benefit implicit memories, which are more easily captured with measures like priming, more than explicit, declarative memories (Henke 2010). These possibilities also fit well with TTT, which posits that differences in how a memory is expressed may be explained both by the relative strength of different neural traces at different time points, and by which of multiple neural traces is most suitable for the demands of the current task. It may be that neural representations supporting implicit integration may require time and consolidation to develop, while neural representations that can leverage explicit retrieval strategies may be available immediately after learning. The use of multiple behavioral tasks that capture both explicit and implicit expressions of memories is a fruitful approach that could be used to better characterize how different memory traces may be differentially expressed over time.

What learning mechanism could enable the integration of overlapping temporal regularities? One candidate is forward prediction: when encountering repeated sequences of events, neural representations of the predicted items are formed and strengthened over subsequent repetitions of the sequence (Kok et al., 2012; Schapiro et al., 2012; Turk-Browne et al., 2012; Schapiro et al., 2013; Hindy et al., 2016; Schapiro et al., 2016; Kok & Turk-Browne, 2018; for a review of hippocampal involvement, see Davachi & DuBrow, 2015). In our case, due to the intermixed exposure to the same AB and two different C objects, forward prediction of both C objects while viewing B may have helped to cement their association. Furthermore, forward prediction may not be the only mechanism that can give rise to the integration, as there is neural evidence that second-order associations built with predictable first-order associations are reflected in patterns of brain activity, suggesting that they can be learned in the absence of predictive mechanisms (Schapiro et al., 2013, 2016). However, the fact that we only observed behavioral integration after a delay suggests that forward prediction may not be sufficient to give rise to behavioral integration across events.

### Cortical and hippocampal similarity

We found that after 24 hours, patterns of activation in mPFC reflected differentiation of objects from overlapping sequences. This was counter to our prediction that with consolidation, neural patterns in mPFC would reflect similarities across sequences with shared temporal regularities. There are a few noteworthy findings of mPFC differentiation that may help to make sense of this finding. First, Ezzyat and colleagues (2018) observed that neural patterns evoked by associative memories in mPFC were more distinct than those evoked by item memories, but this difference only held for events that were re-studied after a night of sleep and not for events re-studied in the same experimental session. Second, individual autobiographical memories can be successfully decoded in mPFC, and classification accuracy for individual memories is greater for more remote memories relative to recent ones (Bonnici et al., 2012). Third, differentiation in mPFC has been shown to emerge over repeated testing, in addition to over a delay period, and also relates to long-term memory (Wirebring et al., 2015), in line with the idea that repeated testing can accelerate the stabilization of memories by minimizing competition with related memories (Antony et al., 2017; Hulbert & Norman, 2015). Taken together, this suggests that mPFC plays a role in the differentiation of discrete memories to support their long-term storage.

Perhaps the most surprising results we found were that despite this *differentiation* in mPFC when considering group-level changes, mPFC neural patterns reflecting *integration* positively scaled with priming across participants. This suggests that the level of neural representational overlap in the same subregions of mPFC may support both differentiation and integration of overlapping memories – extending beyond findings that different subregions within mPFC separately support differentiation and integration (Schlichting et al., 2015). Reports of correlations between neural and behavioral integration despite neural differentiation at the group level have been reported before (Molitor et al., 2020). This suggests that the restructuring of memories in mPFC conforms to ongoing goal states in addition to the properties of the experience itself (e.g. whether there are similarities or differences across events). Consistent with this, in past work, we found that neural pattern overlap increased in mPFC for objects paired with the same scene after a week, but these effects were observed in a source memory test focusing on the scenes, which either overlapped or were distinct from other memories, and less so in a standard recognition test of the experiment-unique objects (Tompary & Davachi, 2017). Further, in a similar experiment where participants were required to make more fine-grained source memory judgements, mPFC patterns only reflected increasing neural integration over time when the encoded object-scene pairs were congruent with prior knowledge (Audrain, 2020). These mixed results suggest that task demands matter: they likely prioritize processes that are most needed for successful performance (Brunec et al., 2020), which could give rise to differentiated or integrated neural patterns evoked by the same stimuli depending on the task. Taken together, however, the extant data combined with past results supports the conclusion that dynamic neural representational change occurs in the mPFC that structures our experiences, but that it may be best to think of these representations as lying on a continuum from separated to integrated, and thus able to support behavior across a variety of tasks.

In this study, the pattern of activity in the anterior hippocampus reflected learned associations between objects that always appeared back-to-back (A and B). This is a direct replication of prior work (Schapiro et al., 2012) and also highlights the importance of hippocampal representations in memory for temporal order (DuBrow & Davachi, 2013; Hsieh et al., 2014; Kalm et al., 2013; Paz et al., 2010). At the same time, there are several reports of hippocampal integration across events that are separated in time, both immediately after learning (Ritchey et al., 2015; Schapiro et al., 2016, p. 201; Schlichting et al., 2015) and after a delay (Dandolo & Schwabe, 2018; Ritchey et al., 2015; Tompary & Davachi, 2017), and even lesion work demonstrating the necessity of the hippocampus for such integration behaviors (Pajkert et al., 2017; Schapiro et al., 2014). Why then did we not observe changes in neural similarity in the hippocampus that reflected overlap across sequences separated in time? One possibility is that any signal reflecting overlap in the hippocampus may have been too subtle to identify in the exposure phase, as participants were not engaging with the objects in a manner that would promote integration processes. In contrast, signals reflecting AB integration may have been strong enough to appear in the absence of an active integration task because AB sequences were viewed twice as many times and were strongly explicitly remembered. Further, as with any null effect, it could be due to low statistical power or low signal to noise ratio, a common issue for MRI investigations of medial temporal lobe structures.

### Post-encoding rest connectivity

A critical component of systems-level consolidation theories is that long-term memories become stabilized through communication between the hippocampus and cortex, resulting in cortical memory traces. Here, we found that learning object sequences resulted in increased rest connectivity between the posterior hippocampus and LOC, a region that codes for object information. This increase in connectivity was related to the extent of recognition priming across participants, suggesting that coordinated post-learning processing in these regions can facilitate the behavioral integration of overlapping memories. This finding extends a growing body of work demonstrating that hippocampal connectivity with stimulus-selective cortex is related to long-term memory retention (Collins & Dickerson, 2018; de Voogd et al., 2016; Keller & Just, 2016; Murty et al., 2017, 2018; Tambini et al., 2010). However, our findings differ from these observations in one critical way: we found that post-learning rest connectivity related to behavioral integration across overlapping memories, rather than retention of discrete memories. To our knowledge, the only one analogous finding has been observed, in a paradigm where post-learning rest connectivity between anterior hippocampus and the fusiform face area was measured after learning premise pairs in an associative inference paradigm (Schlichting & Preston, 2016). Together this suggests that post-learning rest connectivity may not only support the retention of specific episodic memories, but also underpin integration across them. When might consolidation processes support one over the other? One possibility could be that salient features surrounding learning shape consolidation to prioritize goal-relevant information (i.e. Cowan et al., 2021). However, in our study and the one conducted by Schlichting and Preston, the integration tests were a surprise. Without cues about the test, it remains unclear which feature of memory may be prioritized by consolidation mechanisms, or if both are. Future studies should relate learning-related rest connectivity with distinct behavioral measures that capture both retention of individual events and across-event integration.

We also identified a delay-dependent relationship between changes in rest connectivity and cortical integration of objects from overlapping sequences. Specifically, participants with elevated posterior hippocampal-LOC rest connectivity after the remote learning session also exhibited greater neural integration in mPFC for the remotely studied objects. Both measures explained variance in recognition priming across participants, suggesting that they uniquely support delay-dependent behavioral integration. The timing of these two measures may hint at the directionality of their association: the rest scan occurred immediately after learning on Day 1, and the post-learning similarity values were measured on Day 2. This order parallels predictions from systems-level consolidation theories that cortical neural traces are trained by the hippocampus via coordinated processing (McClelland et al., 1995). An intuitive prediction is that coordination between the hippocampus and a particular cortical region would shape similarity in that same region. Indeed, in one study, pattern similarity in mPFC for objects studied in the same temporal context was related to connectivity between the anterior hippocampus and mPFC (Cowan et al., 2020). Relatedly, in past work, we found that post-learning rest connectivity between anterior hippocampus and mPFC was related to the similarity of overlapping memories in anterior hippocampus (Tompary & Davachi, 2017). Both studies identified the same circuit despite differences in the site of representational change and in their experimental designs. However, the current findings suggest that the connection between post-learning connectivity and similarity is more complex, since we found that changes in similarity in mPFC related to changes in hippocampal connectivity with LOC and not mPFC. Further investigation of relationships between cortical similarity and post-learning processing is a promising avenue for constraining theories of systems-level consolidation and understanding the precise circuit interactions unfolding over time that give rise to consolidated memories.

### Future directions and conclusion

There are two major topics that we have so far neglected to mention but are integral to the study of the consolidation and integration of related events. First, new memories are not learned and consolidated in a vacuum. Prior knowledge has long been known to influence the formation of new event memories (Alba & Hasher, 1983; Anderson, 1984; Bartlett, 1932; Bransford & Johnson, 1972). More recently, much work has been devoted to understanding how prior knowledge influences neural processes that support their encoding, long-term storage, and transformation over time (Audrain, 2020; Bellana et al., 2021; Bonasia et al., 2018; Liu et al., 2016, 2018). In our experiment, all object stimuli were real-world objects, thus the random assignments of object-triplet across participants may have incidentally introduce uneven pre-experimental associations among the sequences (e.g., belonging to the same category, or having similar functions, or sharing salient features). This raises interesting questions about how prior knowledge may influence the integration of new related memories in addition to their long-term stabilization, a promising avenue of future work.

Second, TTT is agnostic to whether consolidation mechanisms require the passage of time, or if there are circumstances in which integrated, cortically-based memories can form quickly. There are now many demonstrations that memories can be rapidly consolidated, as characterized by longer retention durations and increased involvement of mPFC: through repeated retrieval (Antony et al., 2017; Ferreira et al., 2018), through ‘fast-mapping’ of novel words onto new concepts (Coutanche & Thompson-Schill, 2015; Sharon et al., 2011); and for events that are consistent with prior knowledge, as discussed above. Furthermore, discrete events can be integrated into more general knowledge, like schemas, immediately after learning (Brown & Evans, 1969; Kumaran et al., 2009; Posner & Keele, 1968; Richter et al., 2019; Tompary et al., 2020; Tompary & Thompson-Schill, 2021; Zeng et al., 2021). However, this does not directly conflict with TTT, as new schemas are increasingly used after a delay even though they are immediately available (Tompary et al., 2020; Zeng et al., 2021), consistent with TTT’s prediction that the form of memory that is expressed (detailed or schematic) is governed by the relative strength of the neural trace of that memory in the hippocampus versus cortex. Future work should test whether the consolidation-related neural measures presented in this study may similarly underpin the rapid integration of new information.

Finally, we note an important caveat. First, to manipulate the passage of time/course of consolidation within participants, we split encoding into two sessions and intermixed encoded memoranda into one test session. There are several advantages and disadvantages to this design over a design in which with one encoding session and retrieval split into ‘immediate’ and ‘delayed’ sessions. In our design, participants are only tested once, preventing rehearsal of stimuli to be tested after a delay after exposure to the test format in the ‘immediate’ session. An intermixed test also minimizes the chances that participants are switching retrieval strategies or response criteria for recent and remotely learned memoranda. However, encoding in the second ‘recent’ session may be influenced by the first ‘remote’ session. Indeed, knowledge of the task is reflected in the overall slower RTs but a larger facilitation in RTs across learning in the remote session relative to the recent one (Supplemental Figure 1A). As we discussed, it is well known that prior knowledge can shape the encoding of new, similar events, often with a shift to circuits involving mPFC when encoding of events that are consistent with prior knowledge (e.g. van Kesteren et al., 2010). This may explain why we find increased rest connectivity of LOC and posterior hippocampus after the first, ‘remote’ session but increased rest connectivity of LOC and mPFC after the second, ‘recent’ session. In summary, neither a design with split encoding nor a design with split retrieval can perfectly isolate influences of consolidation. Ideally, in the future, consolidation effects will be observed in converging findings from both types of designs.

Taken together, the present results demonstrate that over time, episodic memories that share temporal regularities become behaviorally and neurally integrated. These findings provide evidence for the notion that psychological transformations in memories are accompanied by shifts in cortical similarity and driven by post-learning reactivation. These observations reveal new insights into the development of conceptual or semantic memory over time, suggesting that the consolidation of memories with overlapping temporal structures may constitute a key mechanism underlying memory integration.

## Methods

### Experiment 1

#### Subjects

90 right-handed, native English speakers participated in this experiment. Demographic information was lost for 25 participants; of the 65 with intact records, 42 were female, and the mean age was 23 (range: 18-33). Participants were recruited from New York University and the broader community. The University Committee on Activities Involving Human Subjects approved all recruitment and consent protocols. 12 participants were excluded: 6 did not return for the second experimental session; 1 fell asleep; 5 ended early due to experimental error. A further 5 participants were excluded due to poor performance on the recognition test, (A < 3 SD relative to group average for either recent or remote memory), leaving 73 participants included in all following analyses.

#### Stimuli

The stimuli used in this experiment consisted of 100 color images of objects, taken from online databases and used in previous studies (DuBrow & Davachi, 2013, 2014).

#### Experiment procedure

The experiment consisted of two sessions, separated by approximately 24 hours. In the first session, participants incidentally encoded sequences of the objects while performing a cover task (remote learning). In the second session, participants began by incidentally learning a new set of object sequences while performing the same cover task as in the first session (recent learning). Then, participants completed two tests consisting of both sets of stimuli from the two learning sessions: a recognition priming task and an explicit integration task. Because memory for all object sequences was tested at the end of the second scan session, memory for objects learned on the first day were considered remote, and memory for objects learned on the second day were considered recent. Thus we refer to the first session as the *remote* session and the second session as the *recent* session.

#### Sequence learning task

In both sessions, participants incidentally encoded a set of sequences. As a cover task, participants viewed each object and decided if the object present on screen was bigger or smaller than the prior object. All objects were presented as the same size, but participants were instructed to use estimates of their real life sizes.

The order of the objects was arranged such that the first two objects in a sequence (A and B) were presented together 100% of the time. B was directly followed by one of two other objects (C_1_ and C_2_), each following B 50% of the time. Thus, participants were exposed to ABC_1_ half of the time, and ABC_2_ the other half of the time. Six unique sequences of objects were presented in each of the two learning sessions. Six baseline objects were randomly inserted between sequences, for use as control comparisons for C objects (see Methods for priming task below). This resulted in a total of 30 objects in each learning session: 6 A objects, 6 B objects, 6 C_1_ objects, 6 C_2_ objects, and 6 baseline objects.

Each sequence of ABC_1_ and ABC_2_ was repeated 16 times, meaning that A and B were exposed to participants 32 times and C_1_ and C_2_ were exposed to participants 16 times over the course of learning. Baseline objects were presented 16 times to equate exposure frequency with C objects. In other words, ABC_1_ and ABC_2_ sequences would be intermixed either with baseline objects or other sequences. The presentation of all sequences and baseline objects were divided into 4 11-min learning blocks per session, and participants were given the option to take a 1-min break between blocks. The order of the stimuli was pseudo-randomized such that all sequences (ABC_1_ and ABC_2_) appeared four times in each block, all baseline objects appeared twice in each block, and no sequence with the same A and B objects was presented back-to-back.

The presentation timing was designed in anticipation of the planned fMRI experiment (see Experiment 2). Each object was presented for 2-sec, and participants were instructed to respond as quickly as possible before the object was removed from the screen, without sacrificing accuracy. Between each trial, there was a variable inter-trial-interval (ITI) ranging from 0 to 4 seconds. The ITI was constructed to optimize item-level pattern similarity analyses between C_1_ and C_2_ within a sequence (not reported in this manuscript), while also allowing for sufficient jittering of trials to perform condition-level univariate analyses. The ITIs between sequences and before and after baseline objects were randomly jittered.

Finally, the sizes of objects in each sequence were pre-determined such that the motor response to C objects were matched within each sequence: both C_1_ and C_2_ were either bigger or smaller than B. Furthermore, in half of the sequences in each learning session, C_1_ and C_2_ were bigger than B, and in the other half, C_1_ and C_2_ were smaller than B.

#### Recognition priming task

In this task, all 60 objects from both learning sessions were presented intermixed with 40 novel foils. As a cover task, participants were instructed to endorse each object as ‘old’ or ‘new’. They were asked to respond as quickly as possible without sacrificing accuracy, and they were not given an opt-out or ‘don’t know’ option. The ITI between each trial was fixed at 1 second.

Unbeknownst to the participants, the order of the objects was manipulated to use response priming as an index of participant’s implicit knowledge of the association between C_1_ and C_2_. A priming manipulation was used to test memory for ‘across-episode’ associations: namely, the association between C objects that followed the same sequence of A and B. Critically, ABC_1_ and ABC_2_ were never experienced together, but through the overlap in A and B objects, we expect the two episodes to become associated. Note that to participants, C_1_ and C_2_ objects hold an identical mnemonic status, as they both follow B an equal number of times and are presented within their sequence in randomized order over learning. We have assigned them with separate labels for the sole purpose of clarifying the conditions of the priming manipulation. If participants successfully associated C_1_ and C_2_ through their shared prior sequential information, we expected that the presentation of C_1_ before C_2_ would facilitate processing of C_2_, resulting in a faster response time (RT). Each C_1_ object directly preceded the C_2_ object that shared an overlapping sequence of A and B during learning. As a control comparison, each C_1_ object followed a baseline object learned in the same session. Since C and baseline objects share no sequential information, we predicted that the baseline object would not facilitate of processing of C. This order (baseline, C_1_, C_2_) also controlled for motor response history, as we only included objects that (1) were correctly endorsed as ‘old’ and (2) whose prior objects were also correctly endorsed as ‘old’.

We additionally arranged B objects to always follow the A object from the same learning sequence. Like with the C objects, responses times for A and B were only analyzed if both objects were correctly endorsed as ‘old’ and if the prior object was also correctly endorsed. Note however that these comparisons were secondary to our main planned analyses, and due to constraints in randomization, A objects could either be followed by a different old object or a novel foil; thus the motor history for these objects is not controlled for to the same extent as C objects.

#### Explicit memory task

This task followed the recognition priming task. Due to experimental error, two participants did not complete this task, leaving a sample size of 71. In this task, participants viewed pairs of objects and were asked to rate their familiarity from ‘Not familiar’ to ‘Very familiar’. They responded by clicking on an un-numbered sliding scale. Each pair was presented for 6 seconds, separated by a jittered inter-trial interval ranging from .5 – 1.5 sec. Participants viewed 60 pairs: 12 intact AB pairs; 12 re-arranged A and B objects; 12 intact C pairs; 12 re-arranged C objects; and 12 pairs of randomly paired baseline objects. All objects from sequences were viewed twice, once with its corresponding pair and once as a foil in a rearranged pair.

#### Unreported tasks

Participants from Experiment 1 were originally divided into two separate behavioral cohorts that piloted the feasibility of other memory tests that were ultimately not included in Experiment 2. One cohort (N = 48) completed a free recall task, and a second cohort (N = 42) completed a sequence test that mirrored the learning procedure but intermixed sequences from both learning sessions. These tasks were conducted at the end of the second session. Since all aspects of the procedure were identical across the two cohorts until the presentation of those tasks, we have analyzed these two cohorts together to serve as Experiment 1.

#### Statistical Analyses

Linear mixed-effects models were used to quantify recognition priming and to characterize changes in response times and accuracy across learning. Participant intercepts and slope terms for each included predictor variable were modeled as random effects. The significance of a given contrast was obtained using Satterthwaite approximate degrees of freedom, resulting in *F* or *t* statistics and corresponding *p* values. Response times for all learning trials were included in these models, but response times during recognition priming were only included if the current and prior objects were correctly endorsed as ‘old’ (see Recognition Priming Task for more details). Accuracy computed for any learning trial in which the preceding object was coded to be a different size (e.g. an object bigger than a shoebox following an object smaller than a shoebox), as comparisons amongst objects that were both bigger or both smaller are likely more variable and dependent on participants’ subjective opinion.

Explicit memory performance was aggregated for each participant as the average difference in familiarity between Intact and Re-arranged pairs, such that a value of 0 would indicate no reliable discrimination and 1 would indicate maximal discrimination between the two conditions. These values were entered into two-tailed paired t-tests to test for differences in performance between the two learning sessions. One-sample t-tests with μ = 0 were used to test for reliable above-chance performance.

### Experiment 2

#### Subjects

Twenty-eight right-handed, native English speakers (14 female, mean age: 27.14, range: 20-34) participated in this experiment. Participants were recruited from New York University and the broader community. The University Committee on Activities Involving Human Subjects approved all recruitment and consent protocols. Three participants were excluded due to scanner malfunction, and one withdrew after the first session, leaving 24 participants that were included in the following analyses.

#### Procedure

The experiment consisted of two sessions, separated by approximately 24 hours. The encoding and priming tasks were identical to that of Experiment 1, except that encoding was split into 8 scans rather than 4 blocks. Additionally, in the first session, participants also first completed a resting state scan, and then viewed all object stimuli in a random order (pre-learning exposure). Then, participants incidentally encoded sequences of the objects while performing a cover task (remote learning). Finally, participants completed a second resting-state scan. The stimuli were projected onto a screen in the bore of the scanner, and participants viewed them through a mirror attached to the head coil.

In the second session, participants began by incidentally learning a new set of object sequences while performing the same cover task as in the first session (recent learning). After a resting-state scan, participants completed several tasks consisting of both sets of stimuli from the two learning sessions. First, they completed a priming task, and again viewed all stimuli in a random order (post-learning exposure). Finally, participants completed an explicit test of memory integration.

The recognition priming task the only task in the experiment that was not scanned, even though participants completed the task in the scanner between the rest and post-learning similarity scans.

#### Explicit memory task

At the end of the recent session, participants completed two tasks that tested explicit memory for associations between A and B object and associations between C_1_ and C_2_ objects (Figure 2B). In the first scan, participants were presented with an A object at the top of the screen and two B objects at the bottom of the screen. Participants were instructed to choose which B object was paired with the cued A object during learning. The foil B object was from a different sequence in the same learning session. These trials were intermixed with trials where B objects were cued and participants chose the corresponding A object. The same cue/foil structure was used to test explicit memory for the association between C_1_ and C_2_ objects in the second scan. Each trial was presented for 4 seconds with a variable ITI ranging from 3 to 8 seconds.

#### Pre- and post-learning snapshots

Before the remote learning session on Day 1 and after the recent learning session and priming task on Day 2, participants viewed all objects in a pseudo-random order over the course of two 11.8-min scans. The goal of these scans was to create a template pattern for each object stimulus before and after participants learned their temporal associations. With these template patterns, we measured changes in representational similarity between objects that were associated through their temporal structure. Changes in pattern similarity from pre- to post-learning amongst *recently* learned objects would reflect changes in memory representations that corresponded to the temporal structure learned a few minutes beforehand. In contrast, changes in pattern similarity amongst *remotely* learned objects would reflect consolidation-related processes in addition to learning-related changes seen in the recent condition.

Each object was presented for 2 seconds, separated by a variable ITI ranging from 2 to 5 seconds. Participants were instructed to view each object and press a button if a small pound sign (#) appeared anywhere on the image. A pound sign appeared on 20 images (approximately 8% of all trials presented). The objects for which a pound sign appeared were presented a total of five times instead of four, and the trial with the pound sign was omitted from all similarity analyses.

All object stimuli were presented across two scans each in the pre- and post-learning exposure phases. The order during the pre-learning exposure was identical to the order during the post-learning exposure in order to equate any potential confounds due to order effects or biases in modeling the BOLD response. The order of the objects in each scan was randomized for each participant, with the constraint that no object would repeat back-to-back. All A objects and C_1_ objects were presented four times, intermixed in the first scan, and all B objects and C_2_ objects were presented four times, intermixed in the second scan. This arrangement enabled us to prioritize the analysis of similarity between A and B objects, and C_1_ and C_2_ objects, since the objects were presented in separate scans (Mumford et al., 2014).

#### Resting state scans

Participants completed three 6-min resting state scans throughout the experiment: the first and last scans of the remote session, and immediately after sequence learning during the recent session (pre-learning, post-remote learning, post-recent learning). During the scans, participants viewed a blank gray screen and were instructed to remain awake while thinking about whatever they wanted (Greicius et al., 2003; Tambini et al., 2010).

#### fMRI parameters

All scans were collected with a whole-head coil using a 3T Siemens Allegra MRI system. Functional scans consisted of multi-echo gradient-echo planar images (EPI: 2000-ms TR, 15-ms TE, flip angle = 82°, FOV=192×240, 3-mm isotropic voxels), with 34 slices oriented parallel to the AC-PC line. For both sessions, a customized calibration scan was collected using the same slice prescription as the EPI scans. At the end of the second scan, a T1-weighted high-resolution magnetization-prepared rapid-acquisition gradient echo (MPRAGE) sequence (1 x 1 x 1 mm voxels, 176 sagittal slices) was collected.

#### Preprocessing

All learning, rest, and pre- and post-learning exposure scans underwent the same preprocessing steps with FSL (FEAT: http://www.fmrib.ox.ac.uk/fsl). The first six volumes of each EPI were discarded to allow for scanner stabilization. Then, each scan was slice-time corrected and realigned to correct for motion within each run. Smoothing differed based on the analysis: the learning and rest scans were smoothed with a 6mm FWHM Gaussian kernel, while the pre- and post-learning scans were smoothed with a 3mm FWHM Gaussian kernel. All scans were high-pass filtered at 0.01 Hz to remove low-frequency drifts in signal (Cordes et al., 2001) and then aligned to the first pre-learning similarity scan from the remote session.

#### Pre- and post-learning pattern similarity

The four pre- and post-learning scans were entered into separate GLMs after preprocessing. In each scan, a separate 3-sec boxcar regressor was created for the four presentations of each object throughout the scan. One ‘junk’ regressor was created to model the onset of all trials for which participants made a key press. These trials could either be the target trials presented with the pound sign, or any other trial where the participant incorrectly pressed the button in response to a stimulus with no pound sign. As with the learning scans, regressors accounting for head motion were included as well.

These GLMs gave rise to two t-statistic maps for each object presented in the two learning sessions, one from before learning and one from after learning. For each map, the spatial pattern of activity across each ROI was extracted into a vector. These vectors were not z-scored; similarity analyses did not meaningfully change if voxels were z-scored across all patterns. Pearson correlations were computed to measure the neural similarity between vectors of all C objects presented. These were organized into two separate matrices representing C objects from the recent and remote learning sessions. These two matrices were then compared to the neural ‘integration’ model, a matrix of ones and zeros that represented whether C objects were from overlapping or distinct sequences (Figure 3A). A high Point-biserial correlation between neural data and this integration model would reflect greater similarity between images that followed the same AB pairs relative to images that followed different AB pairs. These correlations were Fisher transformed before being entered into statistical tests.

#### Resting state connectivity

The rest scans were used to quantify low-frequency correlations between pairs of ROIs (Albert et al., 2009; Tambini et al., 2010; Tompary et al., 2015), a measure of functional connectivity between regions. To this end, the preprocessed data from the three rest scans were scans were band-pass filtered leaving signal between .01 and .1 Hz (Fox et al., 2005). They were then entered into separate GLMs to model nuisance signals, including: 6 motion regressors and their temporal derivatives; stick functions accounting for sudden head movements; and nuisance signals from white matter tissue and cerebral spinal fluid (CSF) and their temporal derivatives. The motion regressors were derived from frame displacement measurements identifying during motion correction. TRs with sudden head movements were identified with *FSL_motion_outliers*. To create nuisance regressors, each participant’s MPRAGE was segmented into separate masks comprising gray matter, white matter and CSF, using *FAST*. The gray matter and CSF masks were aligned to each participant’s functional volumes and then eroded using *fslmaths*, to minimize the likelihood that these masks contained voxels that partially overlapped with gray matter. Then, the average time course across all voxels in each mask was extracted from the preprocessed rest scans. These time courses were entered into each run’s GLM along with their temporal derivatives.

The residuals of these GLMs were bandpass-filtered, leaving signal ranging from 0.01 and 0.1 Hz, which is the frequency range known to correspond to correlations between gray matter regions in functional neuroimaging data (Cordes et al., 2001). Then, the average time-course for every volume in each rest run was extracted for each ROI. These time courses were then correlated (Pearson correlation), Fisher transformed, and entered into statistical tests.

#### Regions of interest (ROIs)

Bilateral hippocampus was defined using FSL’s automatic subcortical segmentation protocol (FIRST), which anatomically defines subcortical regions using each participant’s T1 anatomical image. To isolate anterior and posterior portions, the number of slices in each hemisphere was divided into three sections, and the anterior and posterior thirds were used as separate ROIs. All ROIs were resampled and aligned to the first similarity scan during the remote session, consistent with the functional scans.

Medial PFC and LOC were functionally defined from the learning scans. Specifically, we sought regions that were sensitive to the transition probabilities of the presented triplets. To identify changes in univariate responses to objects with different transition probabilities, we entered each learning scan into a separate GLM after preprocessing. Two regressors were generated: predictable (B, C_1_, and C_2_), and unpredictable (A, baseline). C_1_ and C_2_ objects were included in the same regressor because participants were unable to differentiate those objects during learning. Trials with no response, or with a response time > 3 SD from the participant’s mean response time, were separately modeled in a regressor of no interest. These regressors and their temporal derivatives were convolved with FSL’s canonical HRF. To account for head motion, the 6 regressors derived from the motion correction procedure were included in each GLM along with their temporal derivatives and stick function regressors derived by *FSL_motion_outliers*.

This resulted in 16 models per participant (8 per session), which were then entered into a fixed-effects analysis in which the first-level estimates were averaged over the two sessions. The resulting contrasts revealed clusters of voxels whose BOLD signal was reliably different for predictable versus unpredictable objects. These estimates were averaged together at the group level in a random-effects analysis. Clusters were determined using a statistical threshold of z > 3.1 and a corrected cluster significance threshold of P < 0.05 using FSL’s Threshold-Free Cluster Enhancement (TFCE).

This analysis revealed several regions including mPFC and LOC (Table 1). As the clusters that overlapped with our target ROIs encompassed other anatomically distinct areas, we masked both ROIs to constrain their coverage. To create the mPFC ROI, we masked the cluster with areas A14m and A10m (Audrain, 2020) from the Brainnettome atlas (https://atlas.brainnetome.org/). To create a bilateral LOC ROI, we masked the two clusters extending over right and left lateral occipital cortex and parahippocampal cortices with the top 90% voxels of the “Lateral Occipital Cortex, Inferior Division” in the Harvard-Oxford Cortical probabilistic atlas.

**Table 1.** R and L indicate right and left hemisphere. Extent is size of the clusters in voxels. x, y, and z coordinates indicate the center of gravity in MNI space (mm). p corresponds to corrected significance of cluster. * indicates clusters used for ROIs.

| Region | Extent | x | y | z | p |
| --- | --- | --- | --- | --- | --- |
| *Predictable > Unpredictable* |  |  |  |  |  |
| * R Parahippocampal, lateral occipital | 4644 | 28.1 | -77.8 | -3.82 | <.001 |
| * L Parahippocampal, lateral occipital | 2623 | -29.1 | -80.3 | -5.08 | <.001 |
| L Angular gyrus, central opercular cortex | 1246 | -52.1 | -43.3 | 27.1 | <.001 |
| * Medial prefrontal cortex, frontal pole | 1195 | -2.43 | 54.5 | 4.31 | <.001 |
| R Angular gyrus, middle temporal gyrus | 911 | 62.2 | -41.2 | 10.1 | <.001 |
| Posterior medial cortex | 675 | -0.133 | -47.5 | 34 | <.001 |
| L. Middle temporal gyrus, posterior | 372 | -56.2 | -24.9 | -5.67 | <.001 |
| Posterior cingulate gyrus | 308 | 1.43 | -21 | 39.6 | <.001 |
| R Parietal operculum cortex | 217 | 59.7 | -24.8 | 21.1 | <.001 |
| R Inferior frontal gyrus | 197 | 55.2 | 31.3 | -1.84 | .001 |
| L Putamen | 168 | -28.5 | -10.8 | 5.01 | .003 |
| L Middle temporal gyrus, anterior | 142 | -51.6 | -1.3 | -23.9 | .008 |
| R Frontal Pole | 98 | 16 | 53.2 | 28.1 | .04 |
| *Unpredictable > Predictable* |  |  |  |  |  |
| L Insula, inferior & middle frontal gyrus | 1788 | -39.7 | 20.1 | 12 | <.001 |
| R Insula, inferior frontal gyrus | 802 | 36.4 | 19.8 | 4.41 | <.001 |
| L Supramarginal gyrus | 252 | -39.4 | -43.7 | 43.1 | <.001 |
| Paracingulate cortex | 221 | -0.922 | 25.8 | 36.4 | <.001 |
| R Middle frontal gyrus | 199 | 44.3 | 33.7 | 21.4 | 0.001 |
| L Inferior temporal gyrus | 142 | -52.7 | -55.2 | -10.6 | 0.008 |
| L Superior lateral occipital cortex | 98 | -22.6 | -65.7 | 38.5 | 0.04 |

#### Statistical tests

Analysis of the behavioral variables was identical to the procedure form Experiment 1, except that explicit memory was aggregated for each participant as the proportion of correct responses. For the neural data, repeated-measures ANOVAs and two-tailed paired t-tests were used to test the significance of group-level effects. Pearson correlations were used to quantify relationships between neural and behavioral measures across participants, and model comparisons were used to test the relative contribution of multiple neural measures in explaining variation in behavior across participants. To examine how rest connectivity and mPFC similarity related to recognition priming (Figure 5B), model comparisons were conducted between nested multiple linear regressions and their significance was determined with *F* values and corresponding *p* values. Results were not corrected for multiple comparisons.

## Supporting information

Supplemental Results

## Acknowledgments

Thank you to Max Bluestone and Yuri Jiao for help with data collection, Emily Cowan for helpful discussions and comments on the manuscript, and Sarah Dubrow for encouragement and mentorship in early stages of the project. The study was supported by National Institute of Mental Health Grant MH074692 (L.D.), Dart Neuroscience (L.D.), and NSF Graduate Research Fellowship Program (A.T.).

## Data availability statement

Raw data and code required to reproduce figures and statistics will be provided for the behavioral data from both experiments. Similarity and connectivity values will be provided with all analysis code needed to reproduce figures and statistics. These materials will be available on OSF upon invitation to submit a Full Submission. Unprocessed brain data will be shared pending publication of this and another manuscript currently in preparation.

## References

1. Acuna, B. D., Eliassen, J. C., Donoghue, J. P., & Sanes, J. N. (2002). Frontal and parietal lobe activation during transitive inference in humans. Cerebral Cortex, 12(12), 1312–1321. https://doi.org/10.1093/cercor/12.12.1312

2. Acuna, B. D., Sanes, J. N., & Donoghue, J. P. (2002). Cognitive mechanisms of transitive inference. Experimental Brain Research, 146(1), 1–10. https://doi.org/10.1007/s00221-002-1092-y

3. Alba, J. W., & Hasher, L. (1983). Is memory schematic? Psychological Bulletin, 93(2), 203–231. https://doi.org/10.1037/0033-2909.93.2.203

4. Anderson, R. C. (1984). Role of the reader’s schema in comprehension, learning, and memory. In: R. Anderson, J. Osborn, & R. Tierney (Eds.), (4th ed.). Newark: International Reading Association. In J. Osborn & R. Tierney (Eds.), Theoretical models and processes of reading (4th ed.). International Reading Association. https://www.oelp.org/wp-content/uploads/2016/03/Role-of-Schemas-Anderson.pdf

5. Antony, J. W., Ferreira, C. S., Norman, K. A., & Wimber, M. (2017). Retrieval as a fast route to memory consolidation. Trends in Cognitive Sciences, 0(0). https://doi.org/10.1016/j.tics.2017.05.001

6. Audrain, S. (2020). Schemas provide a scaffold for neocortical integration at the cost of memory specificity over time. BioRxiv, 42.

7. Bartlett, F. C. (1932). A theory of remembering. In Remembering: A study in experimental and social psychology. Cambridge University Press.

8. Batterink, L. J., & Paller, K. A. (2017). Sleep-based memory processing facilitates grammatical generalization: Evidence from targeted memory reactivation. Brain and Language, 167, 83–93. https://doi.org/10.1016/j.bandl.2015.09.003

9. Bellana, B., Mansour, R., Ladyka-Wojcik, N., Grady, C. L., & Moscovitch, M. (2021). The influence of prior knowledge on the formation of detailed and durable memories. Journal of Memory and Language, 121, 104264. https://doi.org/10.1016/j.jml.2021.104264

10. Bonasia, K., Sekeres, M. J., Gilboa, A., Grady, C. L., Winocur, G., & Moscovitch, M. (2018). Prior knowledge modulates the neural substrates of encoding and retrieving naturalistic events at short and long delays. Neurobiology of Learning and Memory, 153, 26–39. https://doi.org/10.1016/j.nlm.2018.02.017

11. Bonnici, H. M., Chadwick, M. J., Lutti, A., Hassabis, D., Weiskopf, N., & Maguire, E. A. (2012). Detecting representations of recent and remote autobiographical memories in vmPFC and hippocampus. Journal of Neuroscience, 32(47), 16982–16991. https://doi.org/10.1523/JNEUROSCI.2475-12.2012

12. Bransford, J. D., & Johnson, M. K. (1972). Contextual prerequisites for understanding: Some investigations of comprehension and recall. Journal of Verbal Learning and Verbal Behavior, 11(6), 717–726. https://doi.org/10.1016/S0022-5371(72)80006-9

13. Brown, B. R., & Evans, S. H. (1969). Perceptual learning in pattern discrimination tasks with two and three schema categories. Psychonomic Science, 15(2), 101–103. https://doi.org/10.3758/BF03336223

14. Brunec, I. K., Robin, J., Olsen, R. K., Moscovitch, M., & Barense, M. D. (2020). Integration and differentiation of hippocampal memory traces. Neuroscience & Biobehavioral Reviews, 118, 196–208. https://doi.org/10.1016/j.neubiorev.2020.07.024

15. Chatburn, A., Lushington, K., & Kohler, M. J. (2014). Complex associative memory processing and sleep: A systematic review and meta-analysis of behavioural evidence and underlying EEG mechanisms. Neuroscience & Biobehavioral Reviews, 47, 646–655. https://doi.org/10.1016/j.neubiorev.2014.10.018

16. Collins, J. A., & Dickerson, B. C. (2018). Functional connectivity in category-selective brain networks after encoding predicts subsequent memory. Hippocampus, 1–11. https://doi.org/10.1002/hipo.23003

17. Cordes, D., Haughton, V. M., Arfanakis, K., Carew, J. D., Turski, P. A., Moritz, C. H., Quigley, M. A., & Meyerand, M. E. (2001). Frequencies contributing to functional connectivity in the cerebral cortex in “resting-state” data. American Journal of Neuroradiology, 22(7), 1326–1333.

18. Coutanche, M. N., & Thompson-Schill, S. L. (2015). Rapid consolidation of new knowledge in adulthood via fast mapping. Trends in Cognitive Sciences, 19(9), 486–488. https://doi.org/10.1016/j.tics.2015.06.001

19. Cowan, E. T., Liu, A., Henin, S., Kothare, S., Devinsky, O., & Davachi, L. (2020). Sleep Spindles Promote the Restructuring of Memory Representations in Ventromedial Prefrontal Cortex through Enhanced Hippocampal–Cortical Functional Connectivity. Journal of Neuroscience, 40(9), 1909–1919. https://doi.org/10.1523/JNEUROSCI.1946-19.2020

20. Cowan, E. T., Schapiro, A. C., Dunsmoor, J. E., & Murty, V. P. (2021). Memory consolidation as an adaptive process. Psychonomic Bulletin & Review. https://doi.org/10.3758/s13423-021-01978-x

21. Dandolo, L. C., & Schwabe, L. (2018). Time-dependent memory transformation along the hippocampal anterior–posterior axis. Nature Communications, 9(1), 1205. https://doi.org/10.1038/s41467-018-03661-7

22. Davachi, L., & DuBrow, S. (2015). How the hippocampus preserves order: The role of prediction and context. Trends in Cognitive Sciences, 19(2), 92–99. https://doi.org/10.1016/j.tics.2014.12.004

23. de Voogd, L. D., Fernández, G., & Hermans, E. J. (2016). Awake reactivation of emotional memory traces through hippocampal–neocortical interactions. NeuroImage, 134, 563–572. https://doi.org/10.1016/j.neuroimage.2016.04.026

24. DeVito, L. M., Lykken, C., Kanter, B. R., & Eichenbaum, H. (2010). Prefrontal cortex: Role in acquisition of overlapping associations and transitive inference. Learning & Memory, 17(3), 161–167. https://doi.org/10.1101/lm.1685710

25. Djonlagic, I., Rosenfeld, A., Shohamy, D., Myers, C., Gluck, M., & Stickgold, R. (2009). Sleep enhances category learning. Learning & Memory, 16(12), 751–755. https://doi.org/10.1101/lm.1634509

26. DuBrow, S., & Davachi, L. (2013). The influence of context boundaries on memory for the sequential order of events. Journal of Experimental Psychology: General, 142(4), 1277–1286. https://doi.org/10.1037/a0034024

27. DuBrow, S., & Davachi, L. (2014). Temporal memory is shaped by encoding stability and intervening item reactivation. Journal of Neuroscience, 34(42), 13998– 14005. https://doi.org/10.1523/JNEUROSCI.2535-14.2014

28. Durrant, S. J., Cairney, S. A., & Lewis, P. A. (2013). Overnight consolidation aids the transfer of statistical knowledge from the medial temporal lobe to the striatum. Cerebral Cortex, 23(10), 2467–2478. https://doi.org/10.1093/cercor/bhs244

29. Durrant, S. J., Taylor, C., Cairney, S., & Lewis, P. A. (2011). Sleep-dependent consolidation of statistical learning. Neuropsychologia, 49(5), 1322–1331. https://doi.org/10.1016/j.neuropsychologia.2011.02.015

30. Ellenbogen, J. M., Hu, P. T., Payne, J. D., Titone, D., & Walker, M. P. (2007). Human relational memory requires time and sleep. Proceedings of the National Academy of Sciences, 104(18), 7723–7728. https://doi.org/10.1073/pnas.0700094104

31. Ezzyat, Y., Inhoff, M. C., & Davachi, L. (2018). Differentiation of human medial prefrontal cortex activity underlies long-term resistance to forgetting in memory. Journal of Neuroscience, 38(48), 10244–10254. https://doi.org/10.1523/JNEUROSCI.2290-17.2018

32. Ferreira, C. S., Charest, I., & Wimber, M. (2018). Testing the fast consolidation hypothesis of retrieval-mediated learning. BioRxiv, 458687. https://doi.org/10.1101/458687

33. Fox, M. D., Snyder, A. Z., Barch, D. M., Gusnard, D. A., & Raichle, M. E. (2005). Transient BOLD responses at block transitions. NeuroImage, 28(4), 956–966. https://doi.org/10.1016/j.neuroimage.2005.06.025

34. Frankland, P. W., Bontempi, B., Talton, L. E., Kaczmarek, L., & Silva, A. J. (2004). The involvement of the anterior cingulate cortex in remote contextual fear memory. Science (New York, N.Y.), 304(5672), 881–883. https://doi.org/10.1126/science.1094804

35. Graveline, Y. M., & Wamsley, E. J. (2017). The impact of sleep on novel concept learning. Neurobiology of Learning and Memory, 141, 19–26. https://doi.org/10.1016/j.nlm.2017.03.008

36. Greene, A. J., Gross, W. L., Elsinger, C. L., & Rao, S. M. (2006). An fMRI analysis of the human hippocampus: Inference, context, and task awareness. Journal of Cognitive Neuroscience, 18(7), 1156–1173. https://doi.org/10.1162/jocn.2006.18.7.1156

37. Greicius, M. D., Krasnow, B., Reiss, A. L., & Menon, V. (2003). Functional connectivity in the resting brain: A network analysis of the default mode hypothesis. Proceedings of the National Academy of Sciences, 100(1), 253–258. https://doi.org/10.1073/pnas.0135058100

38. Gruber, M. J., Ritchey, M., Wang, S.-F., Doss, M. K., & Ranganath, C. (2016). Post-learning hippocampal dynamics promote preferential retention of rewarding events. Neuron, 0(0). https://doi.org/10.1016/j.neuron.2016.01.017

39. Heckers, S., Zalesak, M., Weiss, A. P., Ditman, T., & Titone, D. (2004). Hippocampal activation during transitive inference in humans. Hippocampus, 14(2), 153–162. https://doi.org/10.1002/hipo.10189

40. Henke, K. (2010). A model for memory systems based on processing modes rather than consciousness. Nature Reviews Neuroscience, 11(7), 523–532. https://doi.org/10.1038/nrn2850

41. Hindy, N. C., Ng, F. Y., & Turk-Browne, N. B. (2016). Linking pattern completion in the hippocampus to predictive coding in visual cortex. Nature Neuroscience, 19(5), 665–667. https://doi.org/10.1038/nn.4284

42. Hsieh, L.-T., Gruber, M. J., Jenkins, L. J., & Ranganath, C. (2014). Hippocampal activity patterns carry information about objects in temporal context. Neuron, 81(5), 1165–1178. https://doi.org/10.1016/j.neuron.2014.01.015

43. Hulbert, J. C., & Norman, K. A. (2015). Neural differentiation tracks improved recall of competing memories following interleaved study and retrieval practice. Cerebral Cortex, 25(10), 3994–4008. https://doi.org/10.1093/cercor/bhu284

44. Kalm, K., Davis, M. H., & Norris, D. (2013). Individual sequence representations in the medial temporal lobe. Journal of Cognitive Neuroscience, 25(7), 1111–1121. https://doi.org/10.1162/jocn_a_00378

45. Keller, T. A., & Just, M. A. (2016). Structural and functional neuroplasticity in human learning of spatial routes. NeuroImage, 125, 256–266. https://doi.org/10.1016/j.neuroimage.2015.10.015

46. Kok, P., Jehee, J. F. M., & de Lange, F. P. (2012). Less is more: Expectation sharpens representations in the primary visual cortex. Neuron, 75(2), 265–270. https://doi.org/10.1016/j.neuron.2012.04.034

47. Kok, P., & Turk-Browne, N. B. (2018). Associative prediction of visual shape in the hippocampus. Journal of Neuroscience, 38(31), 6888–6899. https://doi.org/10.1523/JNEUROSCI.0163-18.2018

48. Koscik, T. R., & Tranel, D. (2012). The human ventromedial prefrontal cortex is critical for transitive inference. Journal of Cognitive Neuroscience, 24(5), 1191–1204. https://doi.org/10.1162/jocn_a_00203

49. Kuhl, B. A., Shah, A. T., DuBrow, S., & Wagner, A. D. (2010). Resistance to forgetting associated with hippocampus-mediated reactivation during new learning. Nature Neuroscience, 13(4), 501–506. https://doi.org/10.1038/nn.2498

50. Kumaran, D., Summerfield, J. J., Hassabis, D., & Maguire, E. A. (2009). Tracking the emergence of conceptual knowledge during human decision making. Neuron, 63(6), 889–901. https://doi.org/10.1016/j.neuron.2009.07.030

51. Lau, H., Tucker, M. A., & Fishbein, W. (2010). Daytime napping: Effects on human direct associative and relational memory. Neurobiology of Learning and Memory, 93(4), 554–560. https://doi.org/10.1016/j.nlm.2010.02.003

52. Lerner, I., & Gluck, M. A. (2019). Sleep and the extraction of hidden regularities: A systematic review and the importance of temporal rules. Sleep Medicine Reviews, 47, 39–50. https://doi.org/10.1016/j.smrv.2019.05.004

53. Liu, Z.-X., Grady, C., & Moscovitch, M. (2016). Effects of prior-knowledge on brain activation and connectivity during associative memory encoding. *Cerebral Cortex*, bhw047. https://doi.org/10.1093/cercor/bhw047

54. Liu, Z.-X., Grady, C., & Moscovitch, M. (2018). The effect of prior knowledge on post-encoding brain connectivity and its relation to subsequent memory. NeuroImage, 167(Supplement C), 211–223. https://doi.org/10.1016/j.neuroimage.2017.11.032

55. Luo, Y., & Zhao, J. (2018). Statistical learning creates novel object associations via transitive relations. Psychological Science, 0956797618762400. https://doi.org/10.1177/0956797618762400

56. McClelland, J. L., McNaughton, B. L., & O’Reilly, R. C. (1995). Why there are complementary learning systems in the hippocampus and neocortex: Insights from the successes and failures of connectionist models of learning and memory. Psychological Review, 102(3), 419–457. https://doi.org/10.1037/0033-295X.102.3.419

57. McNeill, D. (1963). The origin of associations within the same grammatical class. Journal of Verbal Learning and Verbal Behavior, 2(3), 250–262. https://doi.org/10.1016/S0022-5371(63)80091-2

58. Milivojevic, B., Vicente-Grabovetsky, A., & Doeller, C. F. (2015). Insight reconfigures hippocampal-prefrontal memories. Current Biology, 25(7), 821–830. https://doi.org/10.1016/j.cub.2015.01.033

59. Molitor, R. J., Sherrill, K. R., Morton, N. W., Miller, A. A., & Preston, A. R. (2020). Memory reactivation during learning simultaneously promotes dentate gyrus/CA2,3 pattern differentiation and CA1 memory integration. Journal of Neuroscience. https://doi.org/10.1523/JNEUROSCI.0394-20.2020

60. Morrissey, M. D., Insel, N., & Takehara-Nishiuchi, K. (2017). Generalizable knowledge outweighs incidental details in prefrontal ensemble code over time. ELife, 6, e22177. https://doi.org/10.7554/eLife.22177

61. Morton, N. W., Schlichting, M. L., & Preston, A. R. (2020). Representations of common event structure in medial temporal lobe and frontoparietal cortex support efficient inference. Proceedings of the National Academy of Sciences. https://doi.org/10.1073/pnas.1912338117

62. Mumford, J. A., Davis, T., & Poldrack, R. A. (2014). The impact of study design on pattern estimation for single-trial multivariate pattern analysis. NeuroImage, 103, 130–138. https://doi.org/10.1016/j.neuroimage.2014.09.026

63. Murty, V. P., DuBrow, S., & Davachi, L. (2018). Decision-making increases episodic memory via postencoding consolidation. Journal of Cognitive Neuroscience, 1–10. https://doi.org/10.1162/jocn_a_01321

64. Murty, V. P., Tompary, A., Adcock, R. A., & Davachi, L. (2017). Selectivity in postencoding connectivity with high-level visual cortex is associated with reward-motivated memory. Journal of Neuroscience, 37(3), 537–545. https://doi.org/10.1523/JNEUROSCI.4032-15.2017

65. Nadel, L., Samsonovich, A., Ryan, L., & Moscovitch, M. (2000). Multiple trace theory of human memory: Computational, neuroimaging, and neuropsychological results. Hippocampus, 10(4), 352–368. https://doi.org/10.1002/1098-1063(2000)10:4<352::AID-HIPO2>3.0.CO;2-D

66. Pajkert, A., Finke, C., Shing, Y.-L., Hoffmann, M., Sommer, W., Heekeren, H. R., & Ploner, C. J. (2017). Memory integration in humans with hippocampal lesions. Hippocampus, 1–9. https://doi.org/10.1002/hipo.22766

67. Paz, R., Gelbard-Sagiv, H., Mukamel, R., Harel, M., Malach, R., & Fried, I. (2010). A neural substrate in the human hippocampus for linking successive events. Proceedings of the National Academy of Sciences, 107(13), 6046–6051. https://doi.org/10.1073/pnas.0910834107

68. Posner, M. I., & Keele, S. W. (1968). On the genesis of abstract ideas. Journal of Experimental Psychology, 77(3p1), 353. https://doi.org/10.1037/h0025953

69. Preston, A. R., & Eichenbaum, H. (2013). Interplay of hippocampus and prefrontal cortex in memory. Current Biology, 23, R764–R773.

70. Preston, A. R., Shrager, Y., Dudukovic, N. M., & Gabrieli, J. D. E. (2004). Hippocampal contribution to the novel use of relational information in declarative memory. Hippocampus, 14(2), 148–152. https://doi.org/10.1002/hipo.20009

71. Richter, F. R., Bays, P. M., Jeyarathnarajah, P., & Simons, J. S. (2019). Flexible updating of dynamic knowledge structures. Scientific Reports, 9(1), 2272. https://doi.org/10.1038/s41598-019-39468-9

72. Richter, F. R., Chanales, A. J. H., & Kuhl, B. A. (2016). Predicting the integration of overlapping memories by decoding mnemonic processing states during learning. NeuroImage, 124, Part A, 323–335. https://doi.org/10.1016/j.neuroimage.2015.08.051

73. Ritchey, M., Montchal, M. E., Yonelinas, A. P., & Ranganath, C. (2015). Delay-dependent contributions of medial temporal lobe regions to episodic memory retrieval. ELife, 4, e05025. https://doi.org/10.7554/eLife.05025

74. Schapiro, A. C., Gregory, E., Landau, B., McCloskey, M., & Turk-Browne, N. B. (2014). The necessity of the medial-temporal lobe for statistical learning. Journal of Cognitive Neuroscience, 1–12. https://doi.org/10.1162/jocn_a_00578

75. Schapiro, A. C., Kustner, L. V., & Turk-Browne, N. B. (2012). Shaping of object representations in the human medial temporal lobe based on temporal regularities. Current Biology: CB, 1622–1627. https://doi.org/10.1016/j.cub.2012.06.056

76. Schapiro, A. C., McDevitt, E. A., Rogers, T. T., Mednick, S. C., & Norman, K. A. (2018). Human hippocampal replay during rest prioritizes weakly learned information and predicts memory performance. Nature Communications, 9(1), Article 1. https://doi.org/10.1038/s41467-018-06213-1

77. Schapiro, A. C., Rogers, T. T., Cordova, N. I., Turk-Browne, N. B., & Botvinick, M. M. (2013). Neural representations of events arise from temporal community structure. Nature Neuroscience, advance online publication. https://doi.org/10.1038/nn.3331

78. Schapiro, A. C., Turk-Browne, N. B., Norman, K. A., & Botvinick, M. M. (2016). Statistical learning of temporal community structure in the hippocampus. Hippocampus, 26(1), 3–8. https://doi.org/10.1002/hipo.22523

79. Schlichting, M. L., Mumford, J. A., & Preston, A. R. (2015). Learning-related representational changes reveal dissociable integration and separation signatures in the hippocampus and prefrontal cortex. Nature Communications, 25(6), 8151. https://doi.org/10.1038/ncomms9151

80. Schlichting, M. L., & Preston, A. R. (2014). Memory reactivation during rest supports upcoming learning of related content. Proceedings of the National Academy of Sciences, 111(44), 15845–15850. https://doi.org/10.1073/pnas.1404396111

81. Schlichting, M. L., & Preston, A. R. (2016). Hippocampal–medial prefrontal circuit supports memory updating during learning and post-encoding rest. Neurobiology of Learning and Memory, 134(Part A), 91–106. https://doi.org/10.1016/j.nlm.2015.11.005

82. Schuck, N. W., Cai, M. B., Wilson, R. C., & Niv, Y. (2016). Human Orbitofrontal Cortex Represents a Cognitive Map of State Space. Neuron, 91(6), 1402–1412. https://doi.org/10.1016/j.neuron.2016.08.019

83. Sekeres, M. J., Winocur, G., & Moscovitch, M. (2018). The hippocampus and related neocortical structures in memory transformation. Neuroscience Letters, 680, 39–53. https://doi.org/10.1016/j.neulet.2018.05.006

84. Sharon, T., Moscovitch, M., & Gilboa, A. (2011). Rapid neocortical acquisition of long-term arbitrary associations independent of the hippocampus. Proceedings of the National Academy of Sciences, 108(3), 1146–1151. https://doi.org/10.1073/pnas.1005238108

85. Spalding, K. N., Schlichting, M. L., Zeithamova, D., Preston, A. R., Tranel, D., Duff, M. C., & Warren, D. E. (2018). Ventromedial prefrontal cortex is necessary for normal associative inference and memory integration. Journal of Neuroscience, 2501–2517. https://doi.org/10.1523/JNEUROSCI.2501-17.2018

86. Squire, L. R., Cohen, N. J., & Nadel, L. (1984). The medial temporal region and memory consolidation: A new hypothesis. In H. Weingartner & E. Parker (Eds.), Memory consolidation (pp. 185–210). Erlbaum.

87. Sweegers, C. C. G., Takashima, A., Fernández, G., & Talamini, L. M. (2014). Neural mechanisms supporting the extraction of general knowledge across episodic memories. NeuroImage, 87, 138–146. https://doi.org/10.1016/j.neuroimage.2013.10.063

88. Sweegers, C. C. G., & Talamini, L. M. (2014). Generalization from episodic memories across time: A route for semantic knowledge acquisition. Cortex, 59, 49–61. https://doi.org/10.1016/j.cortex.2014.07.006

89. Takashima, A., Nieuwenhuis, I. L. C., Jensen, O., Talamini, L. M., Rijpkema, M., & Fernández, G. (2009). Shift from hippocampal to neocortical centered retrieval network with consolidation. The Journal of Neuroscience, 29(32), 10087–10093. https://doi.org/10.1523/JNEUROSCI.0799-09.2009

90. Takashima, A., Petersson, K. M., Rutters, F., Tendolkar, I., Jensen, O., Zwarts, M. J., McNaughton, B. L., & Fernández, G. (2006). Declarative memory consolidation in humans: A prospective functional magnetic resonance imaging study. Proceedings of the National Academy of Sciences of the United States of America, 103(3), 756–761. https://doi.org/10.1073/pnas.0507774103

91. Takehara-Nishiuchi, K., & McNaughton, B. L. (2008). Spontaneous changes of neocortical code for associative memory during consolidation. Science, 322(5903), 960–963. https://doi.org/10.1126/science.1161299

92. Tambini, A., & D’Esposito, M. (2020). Causal contribution of awake post-encoding processes to episodic memory consolidation. Current Biology. https://doi.org/10.1016/j.cub.2020.06.063

93. Tambini, A., Ketz, N., & Davachi, L. (2010). Enhanced brain correlations during rest are related to memory for recent experiences. Neuron, 65(2), 280–290. https://doi.org/10.1016/j.neuron.2010.01.001

94. Tompary, A., & Davachi, L. (2017). Consolidation promotes the emergence of representational overlap in the hippocampus and medial prefrontal cortex. Neuron, 96(1), 228–241.e5. https://doi.org/10.1016/j.neuron.2017.09.005

95. Tompary, A., Duncan, K., & Davachi, L. (2015). Consolidation of associative and item memory is related to post-encoding functional connectivity between the ventral tegmental area and different medial temporal lobe subregions during an unrelated task. The Journal of Neuroscience, 35(19), 7326–7331. https://doi.org/10.1523/JNEUROSCI.4816-14.2015

96. Tompary, A., & Thompson-Schill, S. L. (2021). Semantic influences on episodic memory distortions. Journal of Experimental Psychology: General, 150(9), 1800–1824. https://doi.org/10.1037/xge0001017

97. Tompary, A., Zhou, W., & Davachi, L. (2020). Schematic memories develop quickly, but are not expressed unless necessary. Scientific Reports, 10(1). https://doi.org/10.1038/s41598-020-73952-x

98. Turk-Browne, N. B., Scholl, B. J., Chun, M. M., & Johnson, M. K. (2009). Neural evidence of statistical learning: Efficient detection of visual regularities without awareness. Journal of Cognitive Neuroscience, 21(10), 1934–1945. https://doi.org/10.1162/jocn.2009.21131

99. Turk-Browne, N. B., Simon, M. G., & Sederberg, P. B. (2012). Scene representations in parahippocampal cortex depend on temporal context. The Journal of Neuroscience: The Official Journal of the Society for Neuroscience, 32(21), 7202–7207. https://doi.org/10.1523/JNEUROSCI.0942-12.2012

100. van Kesteren, M. T. R. van, Fernández, G., Norris, D. G., & Hermans, E. J. (2010). Persistent schema-dependent hippocampal-neocortical connectivity during memory encoding and postencoding rest in humans. Proceedings of the National Academy of Sciences, 107(16), 7550–7555. https://doi.org/10.1073/pnas.0914892107

101. Vilberg, K. L., & Davachi, L. (2013). Perirhinal-hippocampal connectivity during reactivation is a marker for object-based memory consolidation. Neuron, 79(6), 1232–1242. https://doi.org/10.1016/j.neuron.2013.07.013

102. Wagner, U., Gais, S., Haider, H., Verleger, R., & Born, J. (2004). Sleep inspires insight. Nature, 427(6972), 352–355. https://doi.org/10.1038/nature02223

103. Warren, D. E., Jones, S. H., Duff, M. C., & Tranel, D. (2014). False recall is reduced by damage to the ventromedial prefrontal cortex: Implications for understanding the neural correlates of schematic memory. The Journal of Neuroscience, 34(22), 7677–7682. https://doi.org/10.1523/JNEUROSCI.0119-14.2014

104. Werchan, D. M., & Gómez, R. L. (2013). Generalizing memories over time: Sleep and reinforcement facilitate transitive inference. Neurobiology of Learning and Memory, 100, 70–76. https://doi.org/10.1016/j.nlm.2012.12.006

105. Wing, E. A., Geib, B. R., Wang, W.-C., Monge, Z., Davis, S. W., & Cabeza, R. (2020). Cortical Overlap and Cortical-Hippocampal Interactions Predict Subsequent True and False Memory. Journal of Neuroscience, 40(9), 1920–1930. https://doi.org/10.1523/JNEUROSCI.1766-19.2020

106. Winocur, G., Moscovitch, M., & Bontempi, B. (2010). Memory formation and long-term retention in humans and animals: Convergence towards a transformation account of hippocampal–neocortical interactions. Neuropsychologia, 48(8), 2339–2356. https://doi.org/10.1016/j.neuropsychologia.2010.04.016

107. Wirebring, L. K., Wiklund-Hörnqvist, C., Eriksson, J., Andersson, M., Jonsson, B., & Nyberg, L. (2015). Lesser neural pattern similarity across repeated tests is associated with better long-term memory retention. The Journal of Neuroscience, 35(26), 9595–9602. https://doi.org/10.1523/JNEUROSCI.3550-14.2015

108. Woodard, J. L., Seidenberg, M., Nielson, K. A., Miller, S. K., Franczak, M., Antuono, P., Douville, K. L., & Rao, S. M. (2007). Temporally graded activation of neocortical regions in response to memories of different ages. Journal of Cognitive Neuroscience, 19(7), 1113–1124. https://doi.org/10.1162/jocn.2007.19.7.1113

109. Yim, H., Savic, O., Sloutsky, V. M., & Dennis, S. J. (2019). Can paradigmatic relations be learned implicitly? Proceedings of the 41st Annual Conference of the Cognitive Science Society, Austin, TX: Cognitive Science Society., 3389.

110. Zeithamova, D., Dominick, A. L., & Preston, A. R. (2012). Hippocampal and ventral medial prefrontal activation during retrieval-mediated learning supports novel inference. Neuron, 75(1), 168–179. https://doi.org/10.1016/j.neuron.2012.05.010

111. Zeng, T., Tompary, A., Schapiro, A. C., & Thompson-Schill, S. L. (2021). Tracking the relation between gist and item memory over the course of long-term memory consolidation. ELife, 10, e65588. https://doi.org/10.7554/eLife.65588

