## Supplemental Results for "Consolidation-dependent behavioral integration of sequences related to mPFC neural overlap and hippocampal-cortical connectivity"

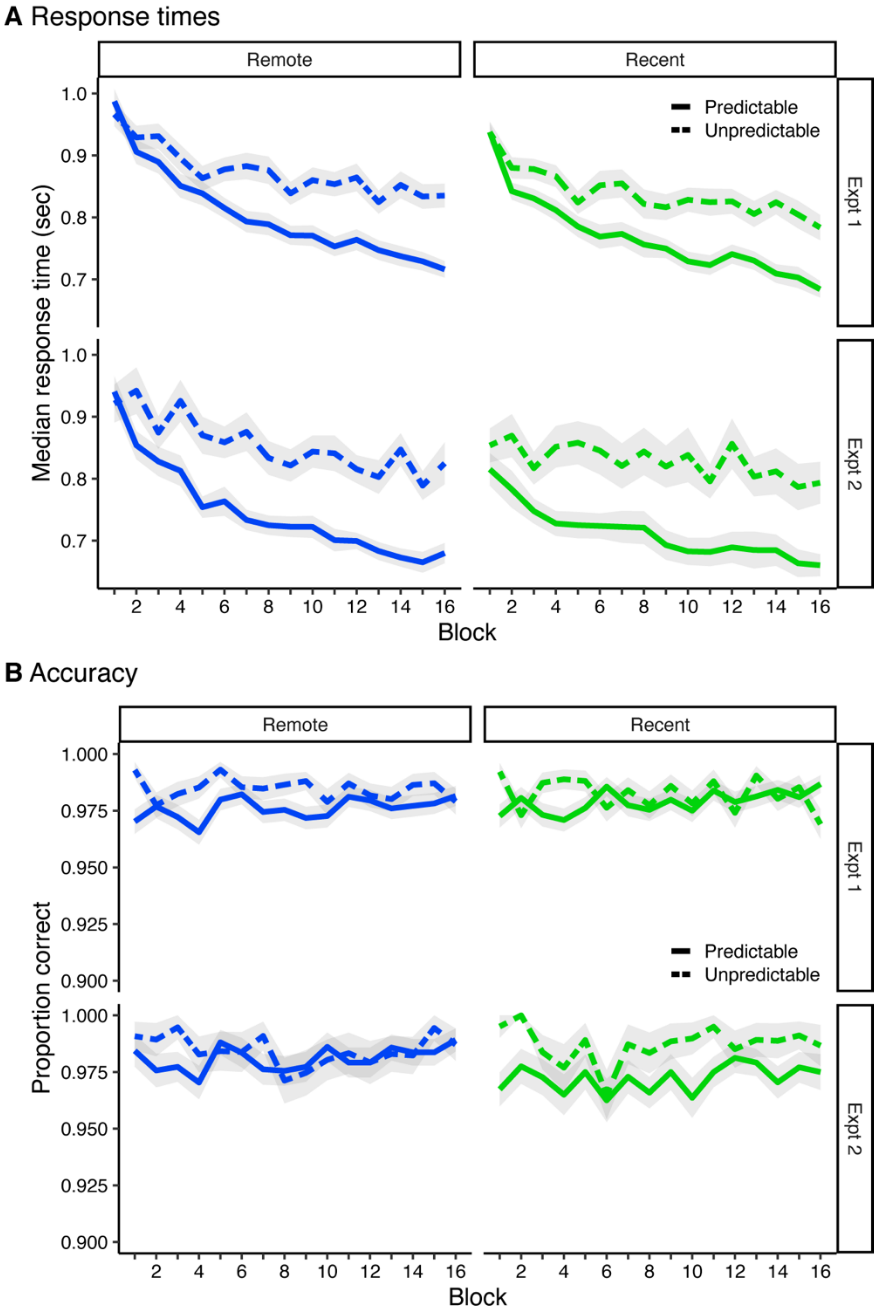


**Supplemental Figure 1, related to Figure 1. (A)** Median response times for predictable items (B and C) and unpredictable items (A and baseline) across learning. Error ribbons signify standard error of the mean (SEM) across participants. **(B)** Average proportion of correct size judgments for predictable items and unpredictable items across learning. Error ribbons signify SEM across participants.

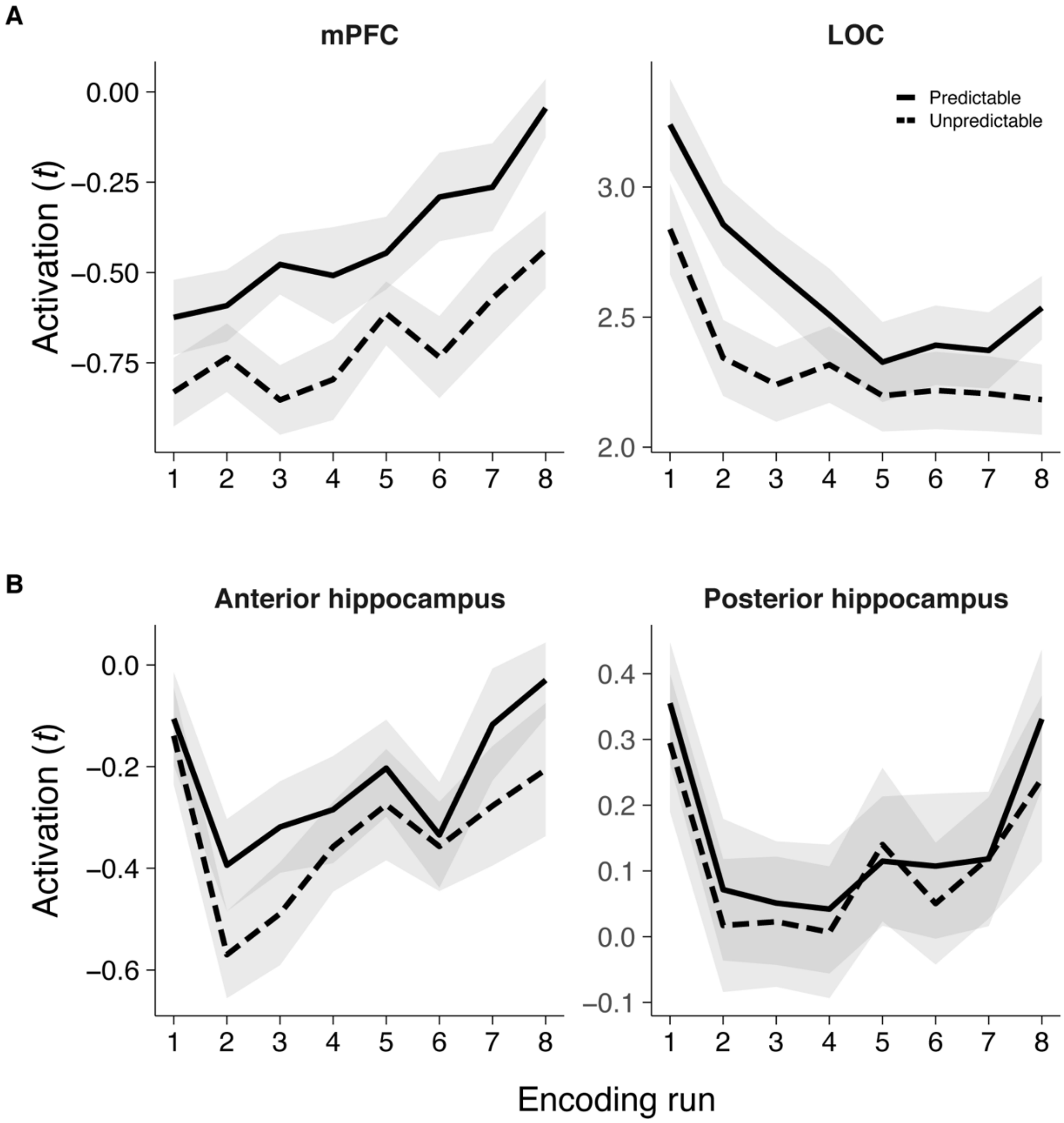


**Supplemental Figure 2, related to Figure 2.** Average univariate activation for predictable and unpredictable items across the eight learning runs, averaged over the recent and remote learning sessions. **(A)** Average univariate activation for mPFC and LOC, both of which were identified from whole brain contrasts comparing predictable and unpredictable items. **(B)** Average univariate activation in anterior and posterior hippocampus, which were anatomically defined. Error ribbons SEM across participants.

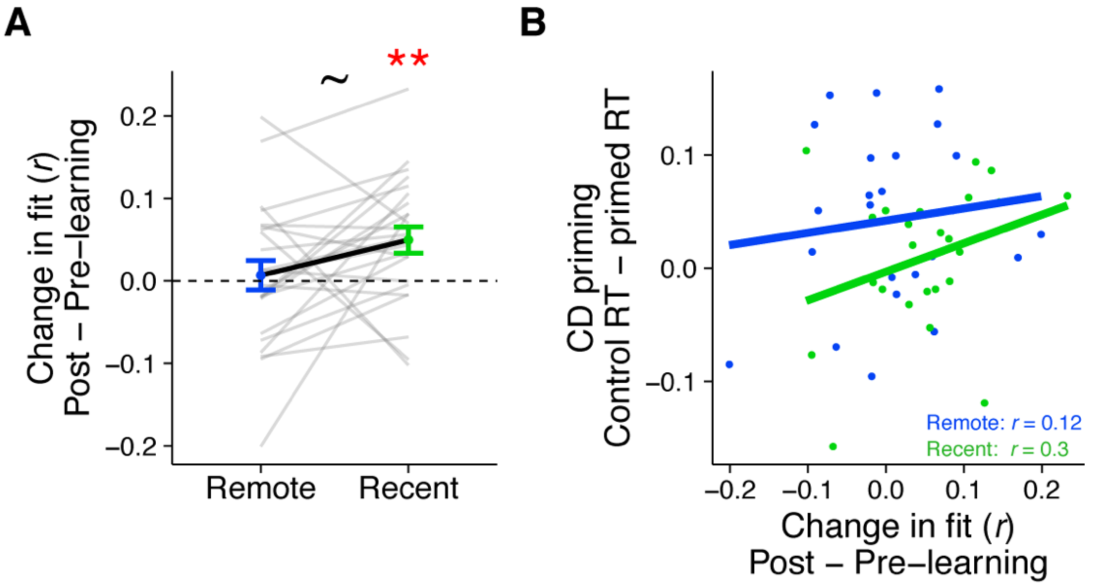


**Supplemental Figure 3, related to Figure 3. (A)** Average change in fit to the neural integration model (post-learning minus pre-learning) in LOC, separately for pairs of C items that were in recently and remotely learned sequences. Values > 0 indicate a better fit to the model after learning, meaning integration of C items from overlapping sequences relative to C items from different sequences. Gray lines indicate participants. Black line indicates group average. Error bars reflect SEM. Black asterisks indicate a two-tailed paired t-test across learning sessions and red asterisks indicate a two-tailed on-sample t-test relative to 0. ~ indicates p < .01; * indicates p < .05. **(B)** Correlation between the average change in LOC similarity and recognition priming across participants, separately for recent and remote learning. Dots indicate participants. Lines indicate line of best fit.

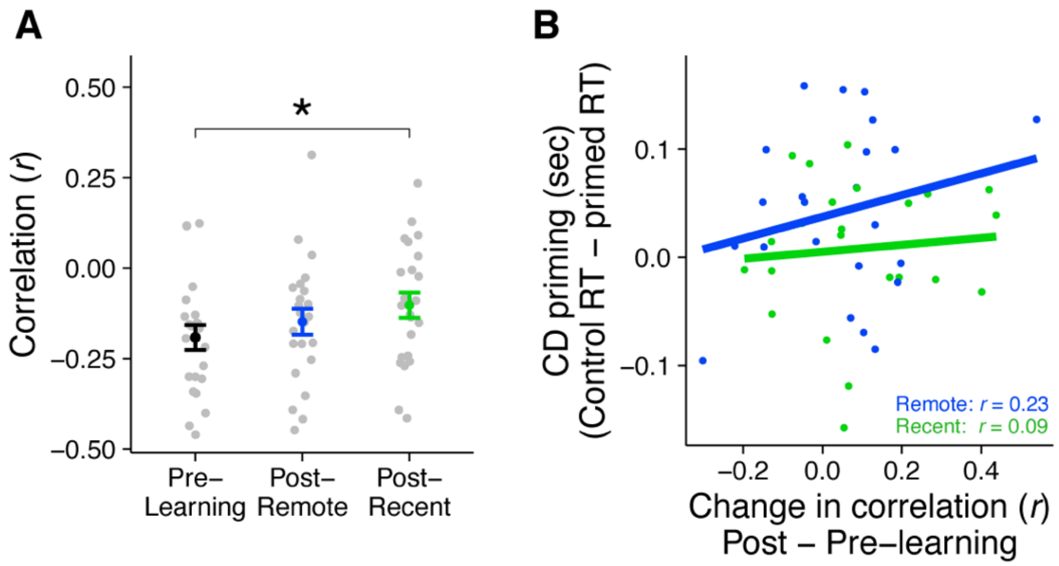


**Supplemental Figure 4, related to Figure 4. (A)** Average rest connectivity between LOC and mPFC. Gray dots indicate participants. Black, blue and green dots indicate group averages. Error bars reflect SEM. * indicates $p < .05$. **(B)** Correlation between the average change in LOC – mPFC connectivity and recognition priming across participants, separately for recent and remote learning. Dots indicate participants. Lines indicate line of best fit.

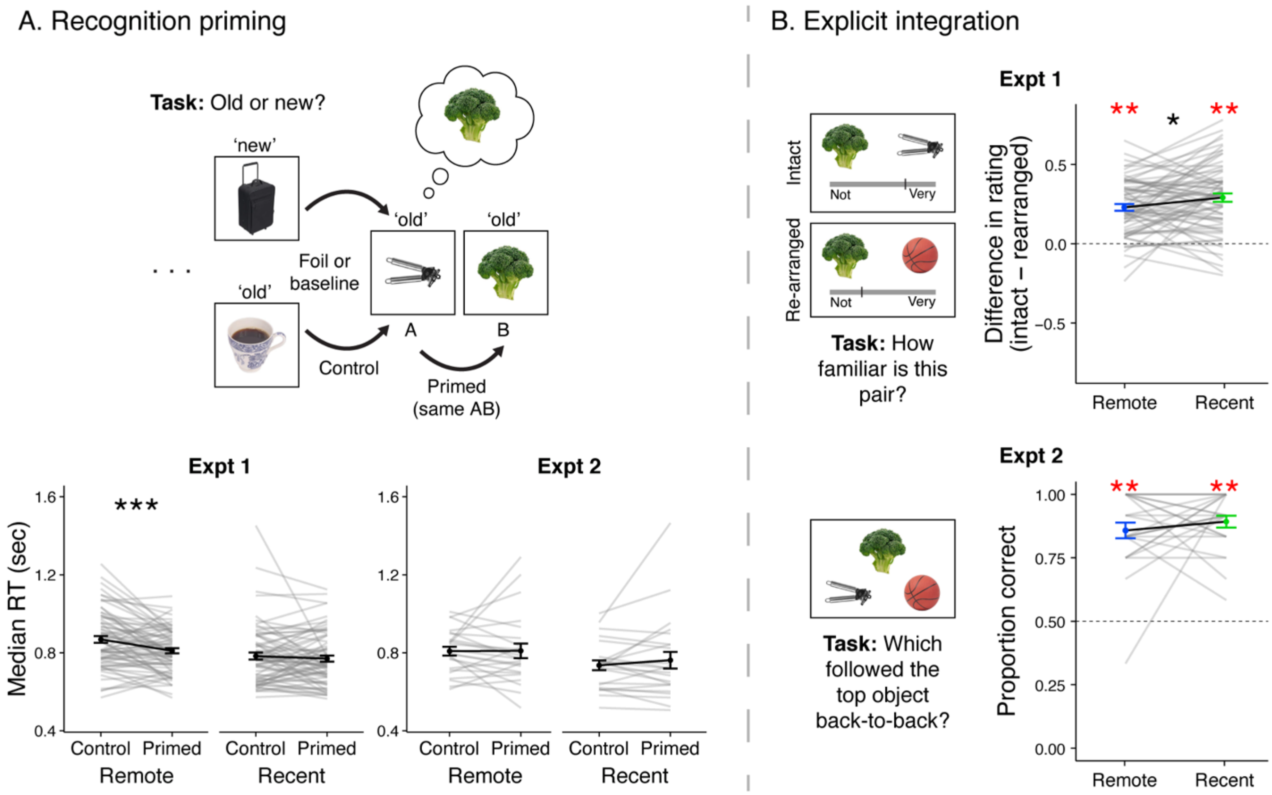


**Supplemental Figure 5, related to Figure 2. (A)** AB priming results. Black lines represent group averages. Gray lines represent participants. Error bars indicate standard error of the mean (SEM) across participants. Statistics reflect paired t-tests across primed and control trials. *** indicates p < .001 **(B)** Explicit integration of AB pairs. Black lines represent group averages. Gray lines represent participants. Error bars indicate standard error of the mean (SEM) across participants. Statistics reflect one-sample t-tests against chance performance. *** indicates p < .001.

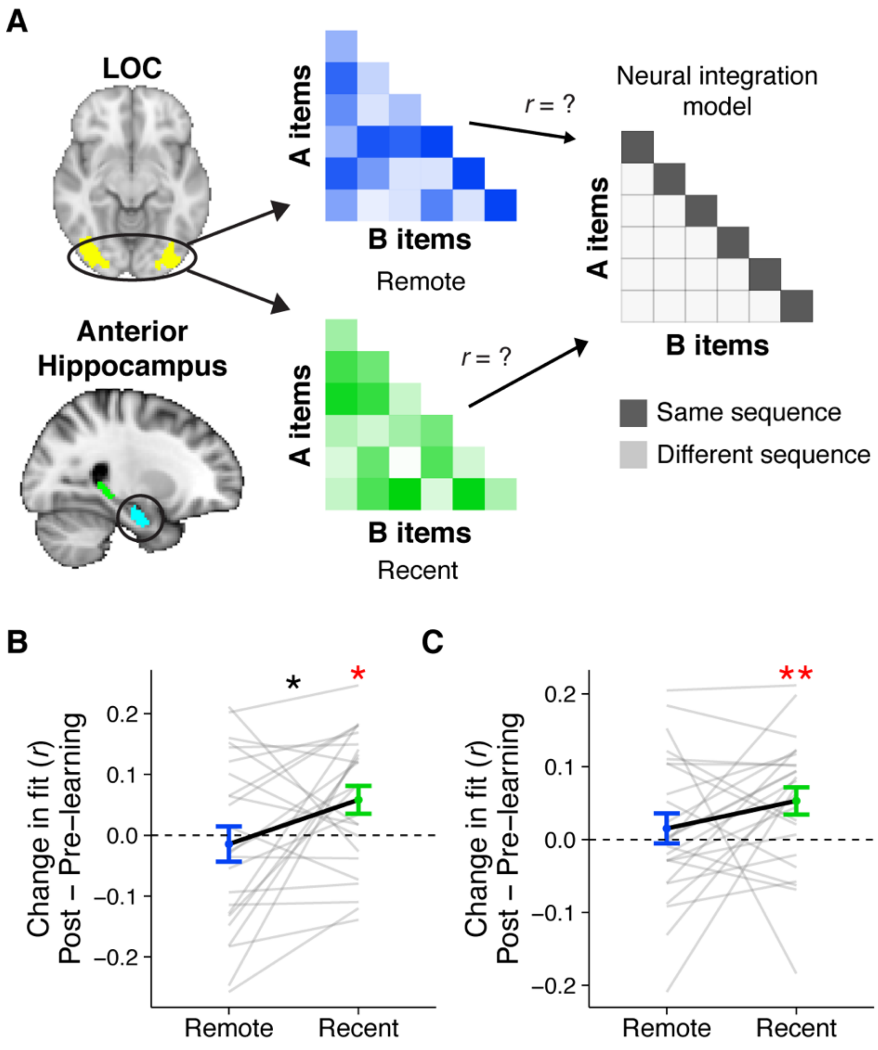


**Supplemental Figure 6, related to Figure 3. (A)** Average change in fit to the neural integration model (post-learning minus pre-learning) in anterior hippocampus and LOC, separately for A and B pairs that were in recently and remotely learned sequences. Values > 0 indicate a better fit to the model after learning, meaning integration of A and B items from overlapping sequences relative to A and B items from different sequences. Gray lines indicate participants. Black line indicates group average. Error bars reflect SEM. Black asterisks indicate a two-tailed paired t-test across learning sessions and red reflects a two-tailed on-sample t-test relative to 0. * indicates $p < .05$.

### Supplemental Results

***Univariate control analyses:*** Here we report several checks to ensure that the observed changes in pattern similarity were not influenced by variability in univariate activation across the different object patterns. First, our pattern similarity approaches relied on point-biserial correlations, which are a special case of Pearson correlations and thus are invariant to average levels of activation across each pattern (meaning that z-scoring across all voxels within a pattern would give rise to the exact correlation matrices as reported in the un-transformed main findings). Second, z-scoring the activation of each voxel across all patterns did not meaningfully change the reported results. Third, we developed two analyses to account for average univariate activation evoked by each item when computing the change in similarity in mPFC and its correlation with recognition priming. Specifically, we generated two additional correlation matrices composed of C1 and C2 items, arranged in the same order as the neural similarity and integration matrices, in order to use them in partial correlations. In the first matrix, for each cell, we computed the average univariate activity between the two items. In the second matrix, we computed the absolute value of the difference in activation across the two items. Both matrices thus represent different possible ways in which the average BOLD signal evoked by two items may be related to their neural similarity – cases where average signal is similarly high or low, or cases where average is close or far apart. We entered each matrix into a separate partial correlation to analyze the change in correlation between the neural similarity in mPFC and the neural integration model, and the extent that this change correlated with recognition priming across participants.

When accounting for the average activation for pairs of items, we found that the change in fit between the correlation matrix reflecting neural similarity and the integration model was significantly negative for the remotely learned items ($t_{(23)} = -3.08$, $p = .005$) and not for recently learned items ($t_{(23)} = -1.38$, $p = .18$), with no reliable difference between the two ($t_{(23)} = -1.41$, $p = .17$). Furthermore, participants with increased similarity amongst C pairs exhibited a stronger priming effect, but only for pairs learned remotely ($r_{(22)} = .58$, $p = 0.003$) and not for pairs learned recently ($r_{(22)} = -.31$, $p = 0.15$).

When accounting for the absolute difference in activation for pairs of items, we found that the correlation between neural similarity and the integration model was significantly negative for the remotely learned items ($t_{(23)}$ = -3.44, p = .002) and not for recently learned items ($t_{(23)}$ = -1.27 p = .22), with no reliable difference between the two ($t_{(23)}$ = -1.60, p = .12). Furthermore, participants with increased similarity amongst C pairs exhibited a stronger priming effect, but only for pairs learned remotely ($r_{(22)}$ = .59, p = 0.003) and not for pairs learned recently ($r_{(22)}$ = -.31, p = 0.14).

In summary, in both control analyses, we replicated the findings observed in the main text: (1) mPFC similarity decreased after learning, particularly after remote learning, and (2) participants with a greater change in similarity in mPFC after learning exhibited stronger recognition priming, only for the remotely learned items. This suggests that the reported findings are not driven by fluctuations in average univariate activation.

***AB integration and similarity:*** In addition to investigating integration across overlapping sequences using the C items, we also included tests of memory for AB pairs, to confirm that participants were able to learn those components of each sequence. We also designed Experiment 2 to replicate findings of integration *within* sequences, in particular the finding that pairs of items that always occur together grow more neurally integrated in the anterior aspect of the hippocampus (Schapiro, Kustner, & Turk-Browne, 2012).

*Recognition:* During the recognition priming task, A and B items were ordered such that all B items followed A items from the same sequence, analogous to the ordering of the C items. We compared response times for B items (primed) to response times to A items, which followed either a novel foil or an item from a different sequence (unprimed). We predicted that responses would be facilitated for primed items over unprimed items, and then this effect would be apparent immediately after learning, like past findings of neural integration of such pairs (Schapiro et al., 2012). Note that due to constraints in pseudo-randomizing the order of these items, the comparison between A and B responses is not well controlled for motor history, unlike the comparison between primed and unprimed C items. For Experiment 1, a trial-level mixed effects linear model was computed with RT as the dependent variable and item (discrete: primed, control), day (discrete: recent, remote) and their interaction as predictors. This model revealed reliable effects of item ($F_{(1,\ 319.18)}$ = 8.30, p = 0.004), and day ($F_{(1,\ 66.53)}$ = 20.65, p < .001), with no reliable interaction ($F_{(1,\ 1332.53)}$ = 2.86, p = .09). Comparisons of response times by item separately for each day revealed that response times were faster for primed items relative to unprimed items for the remotely learned items ($F_{(1,\ 251.0)}$ = 9.82, p = .002) but not the recently learned items ($F_{(1,\ 352.0)}$ = 0.94, p = .33). In Experiment 2, the same day by item model revealed no reliable effect of item ($F_{(1,\ 37.04)}$ = 0.01, p = 0.92), a reliable effect of day ($F_{(1,\ 21.37)}$ = 6.49, p = .02) reflecting slower response times for items from remotely learned sequences, and no interaction ($F_{(1,\ 466.98)}$ = 0.47, p = .50). There were no reliable differences in RT for primed and unprimed items for sequences from either session (both F's < 0.23, both p's > .63). Taken together, participants in Experiment 1 exhibited recognition priming for AB pairs learned 24 hours prior to test, but this effect did not replicate in Experiment 2 (Supplemental Figure 5A).

*Explicit integration:* The same tasks used to assess explicit integration of C items across sequences were also used to assess knowledge of AB pairs. In Experiment 1, participants viewed pairs of A and B items that either came from the same sequence (Intact) or different ones (Rearranged) during learning. They rated the familiarity of these pairs on a sliding scale exactly as for C items. Accuracy was reliably above chance (remote: mean = 0.229, SD = 0.176, $t_{(70)}$ = 11.0, p < .001; recent: mean = 0.291, SD = 0.218, $t_{(70)}$ = 11.2, p < .001) with reliably better memory for pairs from the recent session over the remote session ($t_{(70)}$ = 2.42, p = .02). In Experiment 2, we tested participants' memory for AB pairs with a 2AFC task with either the A or B item as the cue, the other item as the target, and a foil matching the status of the target item (A or B from another sequence). Average accuracy was above chance for items learned remotely (mean = 0.86, SD = 0.152, $t_{(23)}$ = 11.5, p < .001) and recently (mean = 0.892, SD = 0.114, $t_{(23)}$ = 16.9, p < .001) with no difference in accuracy as a function of the two learning sessions ($t_{(23)}$ = -0.84, p = .41). To summarize, in both experiments, explicit memory of AB pairs was quite high, but only in Experiment 1 did accuracy decrease over time (Supplemental Figure 5B).

*Experiment 2 similarity:* To investigate whether neural patterns of A and B items became more integrated over learning and consolidation, we presented all A and B items in the pre- and post-learning exposure scans, intermixed with all C items. We quantified changes in their similarity by extracting patterns of activation for each item and compiling the correlation between all patterns into a correlation matrix with A items in rows and B items in columns. We created separate matrices for items learned in sequences recently or remotely. We then compared each matrix against a model matrix postulating greater similarity between A and B items from overlapping sequences, i.e. items repeated back-to-back 100% of the time, relative to similarity between A and B items presented in different sequences (Supplemental Figure 6A). With this approach, we found increases in AB similarity for recently learned sequences, both in anterior hippocampus (recent: $t_{(23)} = 2.56$, $p = .02$; remote: $t_{(23)} = -0.50$, $p = .62$; Supplemental Figure 6B) and LOC (recent: $t_{(23)} = 2.84$, $p = .009$; remote: $t_{(23)} = 0.74$, $p = .47$; Supplemental Figure 6C). These changes were not related to the extent of recognition priming across sequences (all r's < .25, all p's > .20).
